# Cell fate specification during respiratory development requires ARID1A-containing canonical BAF complex activity

**DOI:** 10.1101/2025.06.02.657302

**Authors:** Hyunwook Lee, Abigail Jaquish, Sharlene Fernandes, Barbara Zhao, Amber Elitz, Kathleen Cook, Sarah Trovillion, Natalia Bottasso-Arias, Simon J. Y. Han, Samantha Goodwin, Nicholas X. Russell, Amanda L. Zacharias, Samantha A. Brugmann, Jeffrey A. Whitsett, Debora Sinner, Xin Sun, Daniel T. Swarr, William J. Zacharias

## Abstract

Development of the mammalian lung requires formation of a definitive lung bud from the foregut endoderm, branching morphogenesis, specification of a proximal to distal axis, and differentiation of extensive specialized airway and alveolar epithelial lineages. These steps require coordinated temporal and regional gene expression to define the lung epithelium in preparation for the first breath at birth. While the transcriptional and signaling regulators required for lung development are increasingly well known, the key epigenetic complexes that interact with lineage transcription factors for cell-specific control of gene expression remain to be defined. Here, we identify a key role for the canonical BAF complex, an ATP-dependent chromatin remodeling complex, during lung epithelial development. Loss of canonical BAF activity throughout the foregut endoderm leads to complete failure of lung formation, and selective deletion of a single key subunit, ARID1A, leads to failure of proximal-distal axis specification, with dilated airway-like structures lined by ectopic basal cells found throughout the distal lung, and failure of specification of alveolar type 1 (AT1) cells in the distal saccular epithelium. In place of AT1 cells, we identified a highly proliferative epithelial cell state defined by joint activation of YAP and WNT signaling and loss of BMP signaling response, leading to increased proliferation and failure of appropriate epithelial differentiation. These changes resulted in secondary failure of mesenchymal and endothelial specification, leading to broad loss of patterning of the distal lung further disrupting peripheral lung morphogenesis. Using embryonic lung organoids, we demonstrate that exogenous BMP4 signaling is sufficient to rescue AT1 and AT2 cell differentiation in ARID1A-null epithelium, while WNT and YAP signaling require functional BAF complex. Together, these data demonstrate a requirement for the BAF complex for lung formation, proximal-distal patterning, and cell fate acquisition, and reveal surprising differential specificity between signaling pathways during lung development.

**One-Sentence Summary:** Functional BAF complex containing ARID1A is required for lung development.

## Introduction

Mammalian lung morphogenesis is marked by a series of coordinated molecular events that begins with the formation of two primary lung buds from the anterior foregut, followed by lung organogenesis. One of the key events during lung development is the establishment and maintenance of the proximal-distal axis. This proximal-distal patterning is tightly controlled by coordinated gene expression changes of signaling molecules and transcription factors that allow for proper development and differentiation of the lung epithelium^1-12^. During distal lung formation, SOX9+/ID2+ progenitor cells in the distal tip epithelium give rise to all cell types of the lung epithelium, though lineage tracing studies show fates become increasingly restricted to alveolar type 1 (AT1) and alveolar type 2 (AT2) epithelial lineages beginning at E13.5^11^. Both proximal-distal axis specification and cell fate allocation require careful balance of proliferation and differentiation by regulating their chromatin state via chromatin remodeling complexes.

One key chromatin remodeling complex is the mammalian SWI/SNF complex, a family of chromatin remodeling complexes possessing ATP-dependent DNA helicase activity that dynamically regulate the accessibility of DNA by sliding and ejecting nucleosomes and by facilitating the writing of H3K27 acetylation marks^13,14^. The canonical BAF complex is one of three distinct known subfamilies of mammalian SWI/SNF complexes that is distinguished by the incorporation of either ARID1A (BAF250A) or ARID1B (BAF250B). ARID1A or ARID1B are required for the complete assembly, stability, and function of the BAF complex^15-17^. ARID1A is required for early embryonic development as ARID1A-null embryos fail to gastrulate^18^. ARID1A contributes to differentiation of stem and progenitor cells in multiple organ systems in both development and adult homeostasis, including several endodermal organs such as the liver, pancreas, and intestines^19-26^. ARID1A is the most frequently mutated gene among the SWI/SNF proteins and BAF complex mutations are associated with human disease^27-29^. Mutations in both ARID1A and ARID1B are found in many forms of malignancy, including non-small cell lung cancer^29,30^. Mutations in ARID1A and ARID1B cause Coffin-Siris syndrome, which is a rare neurological disorder marked by intellectual disability, craniofacial abnormalities, and notably, recurrent pulmonary infections^31,32^. ARID1A and ARID1B mutations are also associated with isolated and complex congenital diaphragmatic hernia presenting with Coffin-Siris syndrome^33,34^. The BAF complex has been extensively studied in these contexts but its role in lung development is unknown.

We recently described a transcriptional regulatory network (TRN) in SOX9+ progenitors at embryonic day 16.5 (E16.5) using scRNAseq and scATACseq which identified the role of the chromatin dynamics during lung morphogenesis^35^. The chromatin regulator most strongly predicted to drive transcription in the E16.5 progenitor TRN was canonical BAF complex, leading us to hypothesize that ARID1A/1B-containing BAF complex may be critical for lung organogenesis. To test this hypothesis, we generated epithelial-specific knockouts of ARID1A and ARID1B using a *Shh*^Cre^ line to ablate canonical BAF complex function throughout the developing lung endoderm. Here, we report that loss of BAF complex activity impairs proximal-distal patterning, distal epithelial fate acquisition, and developmental signaling networks in the embryonic lung.

## Results

### The BAF complex is required in the epithelium for lung morphogenesis

We have previously reported using the paired expression and chromatin accessibility (PECA) model to predict transcriptional regulatory networks (TRN) in E16.5 SOX9^+^ distal lung epithelial progenitors^35^. The PECA model infers TRNs based on localization of cis-regulatory elements, presence of predicted transcription factor binding motifs, and gene expression of putative targets^36^. Based on the predictions of this model, we previously evaluated the role of two transcription factors, PIK3CA and CUX1, in lung development^35,37^. ARID1A and ARID1B were the most highly enriched chromatin regulators in our TRN of developing SOX9^+^ distal epithelial progenitors. Within the network, ARID1A and ARID1B were predicted to interact with other known BAF complex subunits along with several key transcription factors required for lung development (Figure 1A). Predicted targets of the BAF complex in the TRN showed enrichment for a variety of GO Biological Process terms related to cell growth, lung development, and metabolism (Figure 1B). For an unbiased survey of protein-protein interactions of BAF complex in lung epithelium, we performed rapid immunoprecipitation mass spectrometry of endogenous proteins in mouse lung epithelium 15 (MLE-15) cells using ARID1A pulldown, which identified multiple members of the BAF complex, other histone and nucleotide binding proteins (Figure 1C). Together, these data suggested that the BAF complex is a critical regulator of lung development. To test this hypothesis, we deleted both ARID1A and ARID1B using the *Shh*^Cre^ mouse line to generate *Shh*^Cre^;*Arid1a*^fl/fl^;*Arid1b*^fl/fl^ (DKO) embryos to induce a complete loss of canonical BAF complex activity in the respiratory epithelium early in lung morphogenesis (Figure 1D). scRNAseq of control and DKO foreguts at E10.5 showed a dramatic reduction in identifiable lung within the foregut endoderm (Figure 1E). DKO embryos concordantly displayed severe pulmonary hypoplasia at E11.5 and a single tracheoesophageal tube without recognizable lung tissue by E16.5 (Figure 1F-K). Whole-mount immunostaining revealed a near complete loss of NKX2-1^+^ lung epithelium in primordial lung buds and failure of branching morphogenesis (Figure 1L-Q). These data demonstrated that canonical BAF complex activity is required in the respiratory epithelium for lung morphogenesis.

**Figure 1.**
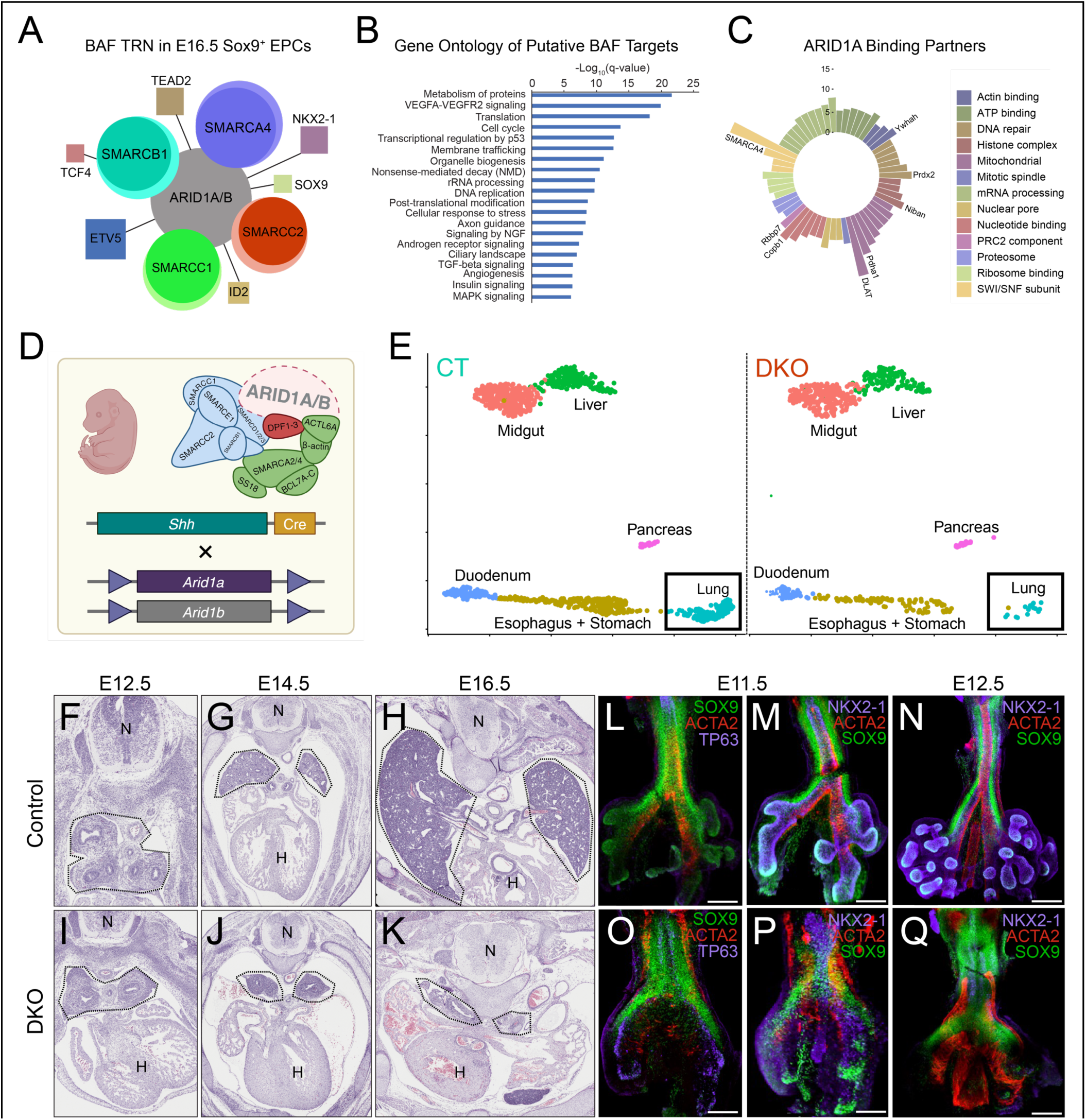
The BAF complex is required in the epithelium for lung morphogenesis. **(A)** Diagram showing transcriptional regulatory network (TRN) for E16.5 SOX9^+^ distal lung progenitors predicted to be regulated by ARID1A and ARID1B. Size of outer circle scaled for number of predicted targets of each component of the BAF complex, size of inner circle scaled by number of targets in common with ARID1A/B. Size of boxes for predicted cooperative TFs scaled by strength of predicted co-regulation with ARID1A/B. (**B)** GO Biological Process term enrichment of putative BAF targets identified in TRN as determined by ToppGene. Enrichments are presented as -log of the corrected p-value. (**C)** Rapid immunoprecipitation mass spectrometry of endogenous proteins (RIME) identifying putative binding partners of ARID1A in MLE-15 cells. **(D)** Experimental design of genetic ablation of ARID1A and ARID1B in the embryonic respiratory epithelium. **(E)** UMAP projection of endodermal clusters from single-cell RNA sequencing of control and DKO embryos at E10.5. Box shows putative lung, which is reduced in DKO embryos. **(F-K)** H&E of formalin-fixed paraffin-embedded tissue sections of control and DKO mouse embryos. **(L-Q)** Whole-mount immunofluorescence of Control and DKO embryos.

### Deletion of ARID1A causes perinatal lethality and aberrant lung morphogenesis

To investigate the relative contribution of ARID1A and ARID1B within the BAF complex, we deleted each individually in the embryonic respiratory epithelium using *Shh*^Cre^ (Figure 2A). While normal Mendelian ratios for control and *Arid1b*^fl/fl^;*Shh*^Cre^ (ARID1B^ShhCre^) mice were observed at P0, 100% of *Arid1a*^fl/fl^;*Shh*^Cre^ (ARID1A^ShhCre^) mice died from respiratory failure at birth (Figure 2B). Histological evaluation across a developmental time series starting from the pseudoglandular stage (E12.5) to the saccular stage of lung development (E18.5) demonstrated no discernable embryonic phenotype or delay in development in ARID1B^ShhCre^ mice (Figure 2C-N). However, ARID1A^ShhCre^ lungs exhibited large, dilated airway-like structures disrupting normal saccular tissue (Figure 2I-J). Therefore, we concluded that ARID1A is required in the BAF complex to drive normal patterning throughout embryonic lung development.

**Figure 2.**
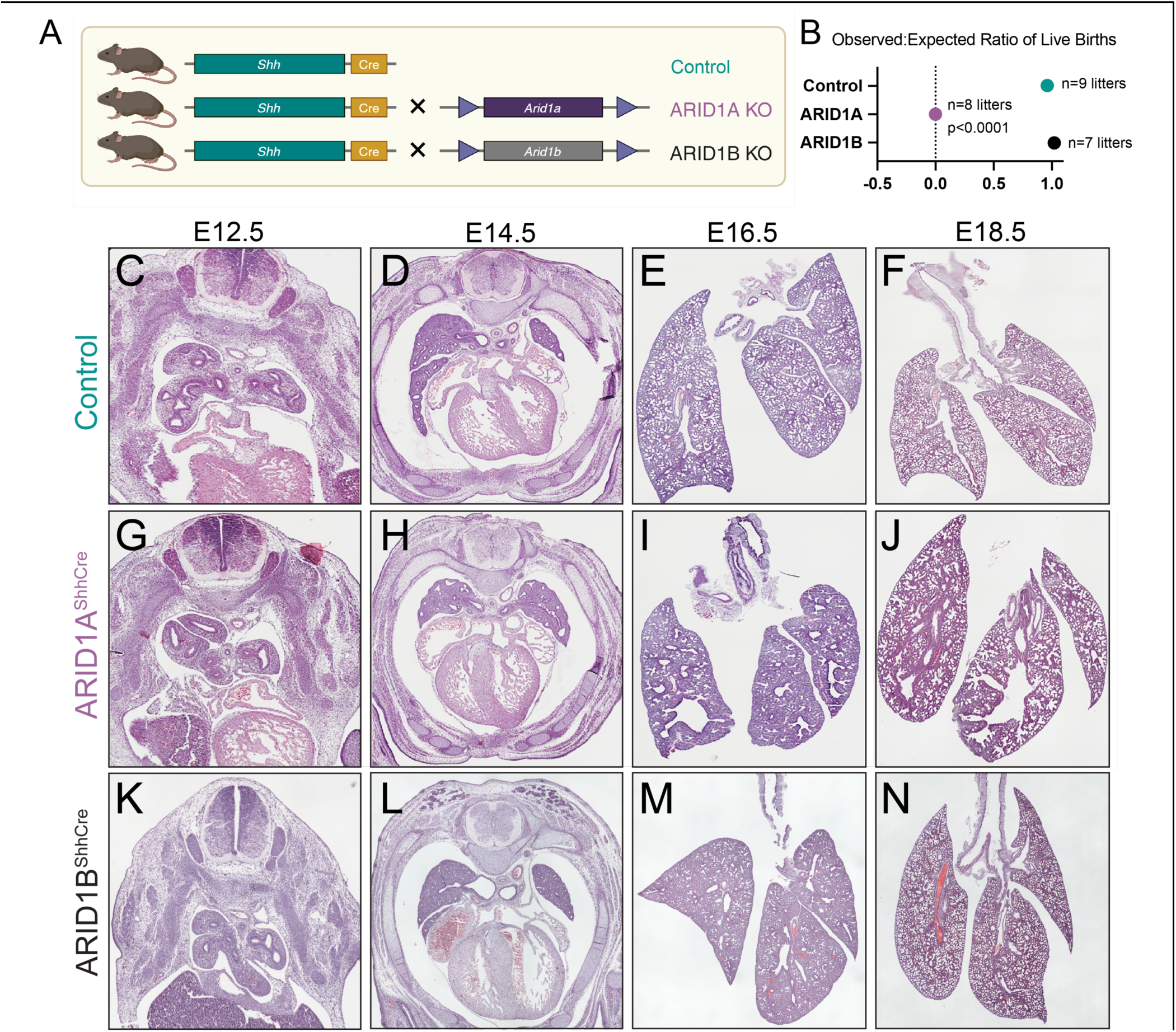
Deletion of ARID1A in the embryonic respiratory epithelium causes in perinatal lethality and aberrant lung morphogenesis. **(A)** Experimental design of genetic ablation of ARID1A or ARID1B alone in the embryonic respiratory epithelium. (**B)** Observed-to-expected ratios of Control, ARID1A^ShhCre^, and ARID1B^ShhCre^ litters. (**C-N)** H&E of formalin-fixed paraffin-embedded tissue sections of Control **(C-F)**, ARID1A^ShhCre^ (G-J), and ARID1B^ShhCre^ **(K-N)** mouse embryos.

### Deletion of ARID1A in the embryonic respiratory epithelium disrupts proximal-distal axis specification

We then turned our attention to detailed evaluation of ARID1A^ShhCre^ mice (Figure 3A). To evaluate the cellular components of the large airway-like structures in ARID1A^ShhCre^ lungs, we performed immunostaining for multiple airway cell type markers. The dilated airway-like structures in ARID1A^ShhCre^ lungs were entirely lined by SOX2^+^ airway epithelial cells (Figure 3B-C), indicating extension of airway-like cells in place of normal saccular tissue in the periphery of the lung. Quantification of SOX2^+^ cells showed that there was a significant increase in airway epithelial cells in the ARID1A^ShhCre^ lung (Figure 3D). scRNAseq of lungs from ARID1A^ShhCre^ mice at E18.5 revealed a concordant increase in airway epithelial cells in mutant embryos. Annotation with LungMAP CellRef identities demonstrated increases in ciliated, deuterosomal, and secretory cell populations. The largest percentage increase was noted in cells annotated as basal cells, which are rarely found in intrapulmonary airways in control mice (Figure 3E-F). Immunostaining for ciliated (TUB1A1), secretory (SCGB1A1), and basal (TP63, KRT5) markers showed extension of ciliated and secretory cells into dilated airway structures in the distal lung (Figure 3D-E, H-I) lined by ectopic basal cells (Figure 3F-G, J-K). P63^+^/KRT5^+^ basal cells were present in the larger intrapulmonary airways of ARID1A^ShhCre^ mice (Figure 3G), and P63^+^/KRT5^-^ “basal-like” cells were present in non-airway structures and in saccular regions of the embryonic lung (Figure 3I/K, O-Q). Since P63 is known to be expressed in early esophageal development in the epithelium, we confirmed that all ectopic P63+ cells expressed the lung lineage marker NKX2-1, consistent with retention of lung rather than gastrointestinal fate. All basal cells in the ARID1A^ShhCre^ distal airways were SOX2^+^ and a subset of this population of cells were also SOX9^+^. SOX2/SOX9 co-staining demonstrated a substantial increase in the number of double SOX9+/SOX2+ in ARID1A^ShhCre^ animals, different from the typical sharp border between SOX2+ airway cells and SOX9+ distal cells present in control E18.5 lungs. Interestingly, “basal-like” cells located in saccular tissue were SOX9^+^ but not SOX2^+^ indicating a partial acquisition of distal fate in these cells despite P63 expression. Together, these data supported the conclusion that ARID1A-containing BAF complex activity is required for proper specification of the proximal-distal axis during lung epithelial differentiation.

**Figure 3.**
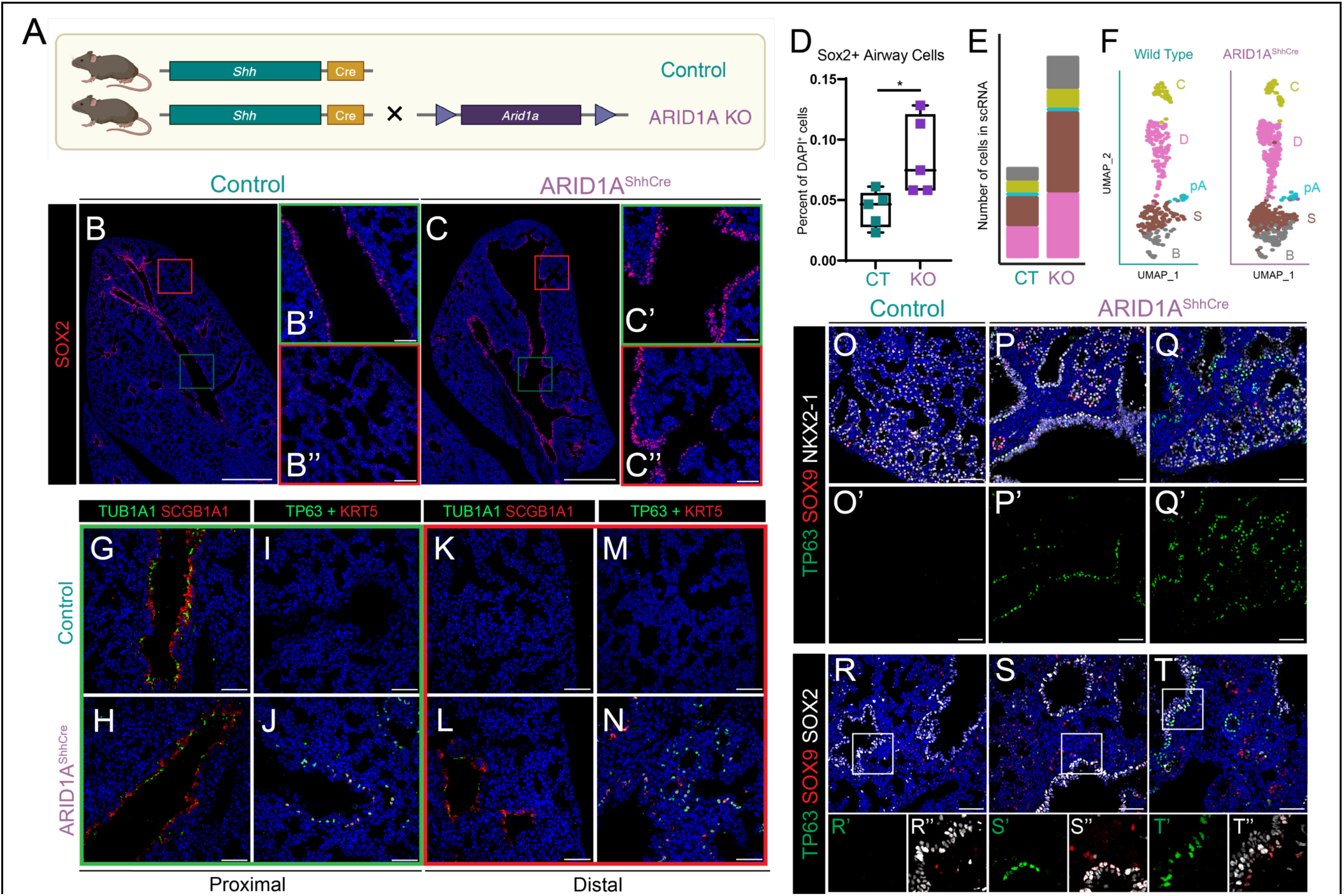
Deletion of ARID1A in the embryonic respiratory epithelium causes failure of the proximal-distal axis specification. **(A)** Design for ablation of ARID1A in the embryonic respiratory epithelium. (**B-D)** Immunostaining for the airway epithelial cell marker SOX2 in central (green box) and peripheral (red box) regions of the lung at E18.5; quantified in (D). **(E-F)** Single-cell RNA sequencing showing increase in number of ARID1A^ShhCre^ airway cell populations at E18.5. **(G-N)** Immunostaining of ciliated (TUB1A1), secretory, (SCGB1A1) and basal cell (KRT5, TP63) markers in Control and ARID1A^ShhCre^ lungs at E18.5 showing both proximal (green box) and distal (red box) regions of lung. **(L)** Quantification of increase in SOX2-positive epithelial cells in ARID1A^ShhCre^ lungs. **(O-Q)** Immunostaining of basal cells (TP63), the distal epithelial progenitor marker (SOX9), and lung epithelial lineage marker (NKX2-1). **(R-T)**. Immunostaining of basal cells (TRP63), the distal epithelial progenitor marker (SOX9), and airway epithelial cell marker (SOX2). Scale bars, 50 μm.

### Airway-specific loss of ARID1A results in normal lung development

Because *Shh* is expressed at the very start of lung morphogenesis before the delineation of the proximal and distal epithelial compartments, we tested whether the aberrant lung morphogenesis seen in ARID1A^ShhCre^ mice was caused by a failure of proximal or distal epithelial fate specification. To specifically ablate ARID1A in the proximal epithelium, we generated *Arid1a*^fl/fl^;*Sox2*^CreER^ (ARID1A^Sox2CreER^) mice, administered tamoxifen orally to pregnant dams at E12.5, and harvested lungs at E18.5 (Figure 4A-B). ARID1A^Sox2CreER^ mice displayed no gross respiratory phenotype at E18.5 (Figure 4C-D). Immunostaining confirmed deletion of ARID1A in the airway epithelium (Figure 4G-H). Both control and ARID1A^Sox2CreER^ mice displayed normal patterning of basal cells in the trachea (Figure 4G-H). While we detected very rare ectopic P63^+^/KRT5^+^ basal cells in some distal airways of ARID1A^Sox2CreER^ mice (Figure 4I-J), the airways were largely indistinguishable from control. Normal patterning of both alveolar type 1 (AT1) and alveolar type 2 (AT2) were also observed in both control and ARID1A^Sox2CreER^ lungs (Figure 4K-R). Thus, the defects observed in lung formation in ARID1A^ShhCre^ mice are likely driven by a failure of distal patterning of ARID1A-null lung epithelium rather than an expansion of ARID1A-null proximal cells (Figure 4S).

**Figure 4.**
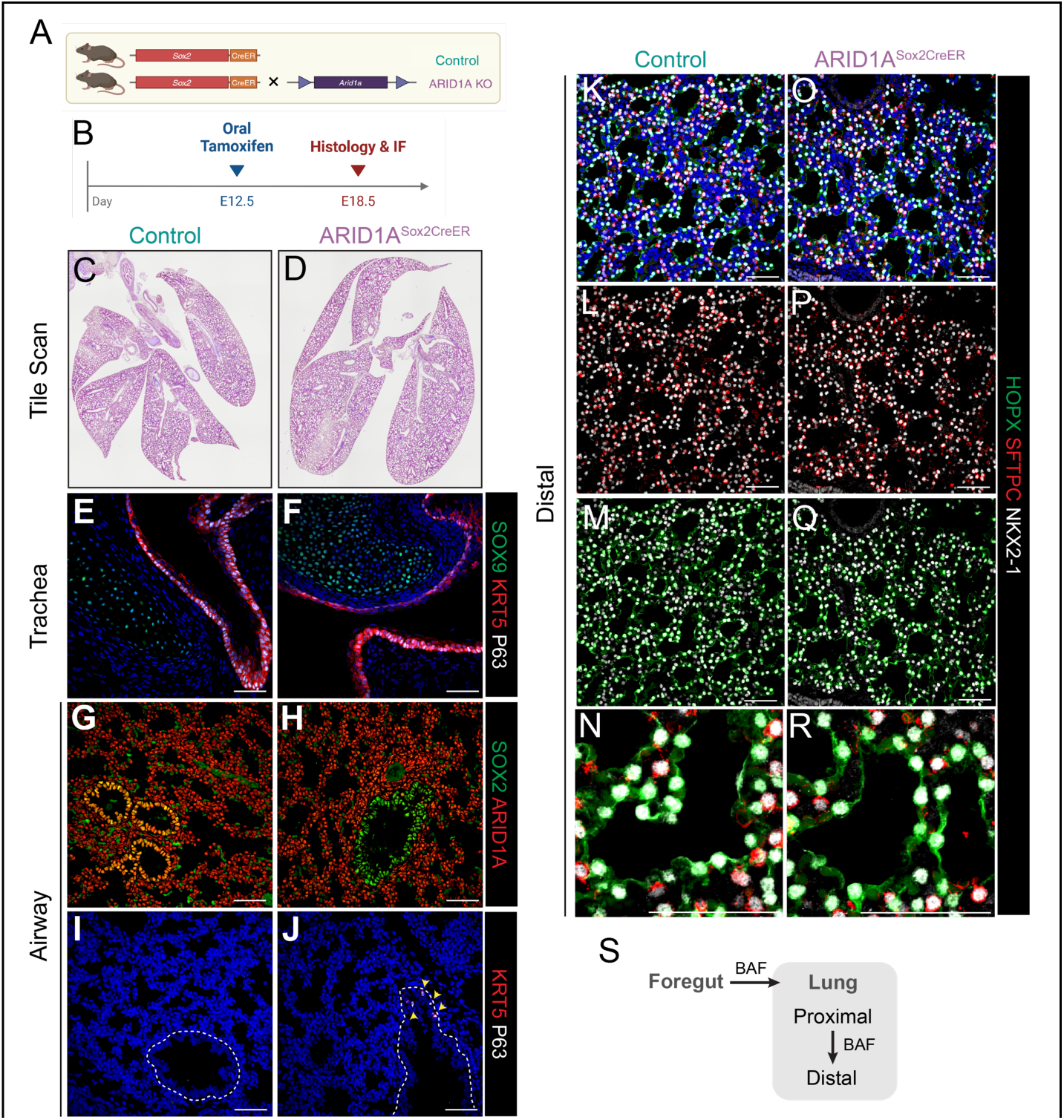
Airway-specific loss of ARID1A results in normal lung development. at E18.5. (**A-B**) Genetic ablation of ARID1A in the embryonic airway epithelium. **(C-D)** H&E stains of Control and ARID1A^SoxCreER^ mice. **(E-F)** Immunostaining for basal cell (KRT5, P63) and chondrocyte (SOX9) markers in CONTROL and ARID1A^SoxCreER^ trachea **(G-J)** Immunostaining for SOX2, ARID1A, KRT5, P63 in CONTROL and ARID1A^SoxCreER^ intrapulmonary airways at E18.5. (**K-R)** Immunostaining of AT1 (HOPX), AT2 (SFTPC), and the lung epithelial lineage (NKX2-1) markers in Control and ARID1A^SoxCreER^ lungs at E18.5. **(S)** Model of BAF function in proximal/distal patterning. Scale bars, 50 μm.

### ARID1A-containing BAF complex is required for distal lung epithelial cell specification

We then turned our attention towards the effect of the loss of epithelial ARID1A on the developing distal saccules. We noted a general decrease in the maturity of saccular structures, with thickened saccular walls and decreased airspace in ARID1A^ShhCre^ mice (Figure 5A-B). Within saccular epithelium, we noted decreased AT1 cells marked by HOPX (Figure 5A-B). Quantification of AT1 and AT2 cell numbers demonstrated a decrease in HOPX^+^/NKX2-1^+^ nuclei in ARID1A^ShhCre^ mice (Figure 5E), and a small but statistically significant decrease in SFTPC+ AT2 cells (Figure 5F). We noted a concomitant increase in HOPX^-^/SFTPC^-^ /NKX2-1^+^ non-AT1 or AT2 cells in saccular locations in ARID1A^ShhCre^ mice (Figure 5G). To investigate the changes in cell populations in the developing saccules at a higher resolution, we evaluated the epithelium from scRNAseq analysis of control and ARID1A^ShhCre^ lungs. Within the saccular epithelium, populations of AT1, AT1/AT2, and Sox9+ distal lung progenitors were reduced in the ARID1A^ShhCre^ lungs compared to the control lungs. Instead, a distinct population with a signature of high proliferation (with expression of *Mki67*) and *Sox9* expression that was transcriptionally unique compared to the SOX9^+^ distal epithelial progenitor population in the control lungs was present in ARID1A^ShhCre^ lungs, which we denote as KO (Figure 5K-L). Immunostaining (Figure 5C-D) and quantification (Figure 5H-J) demonstrated a broad increase in proliferation among NKX2-1^+^ lung epithelial cells (Figure 5H) and an increase in proliferative SOX9^+^ distal epithelial cells (Figure 5I-J) in ARID1A^ShhCre^ mice. GO Biological Process terms associated marker expression within these proliferative ARID1A^ShhCre^-exclusive cells included terms associated with lung development, epithelial differentiation, and sterol and cholesterol biosynthetic processes (Figure 5M). These findings support the conclusion that loss of ARID1A-containing BAF complex activity disrupts distal epithelial differentiation leading to an aberrant population of proliferative SOX9^+^ cells in the distal lung (Figure 5N).

**Figure 5.**
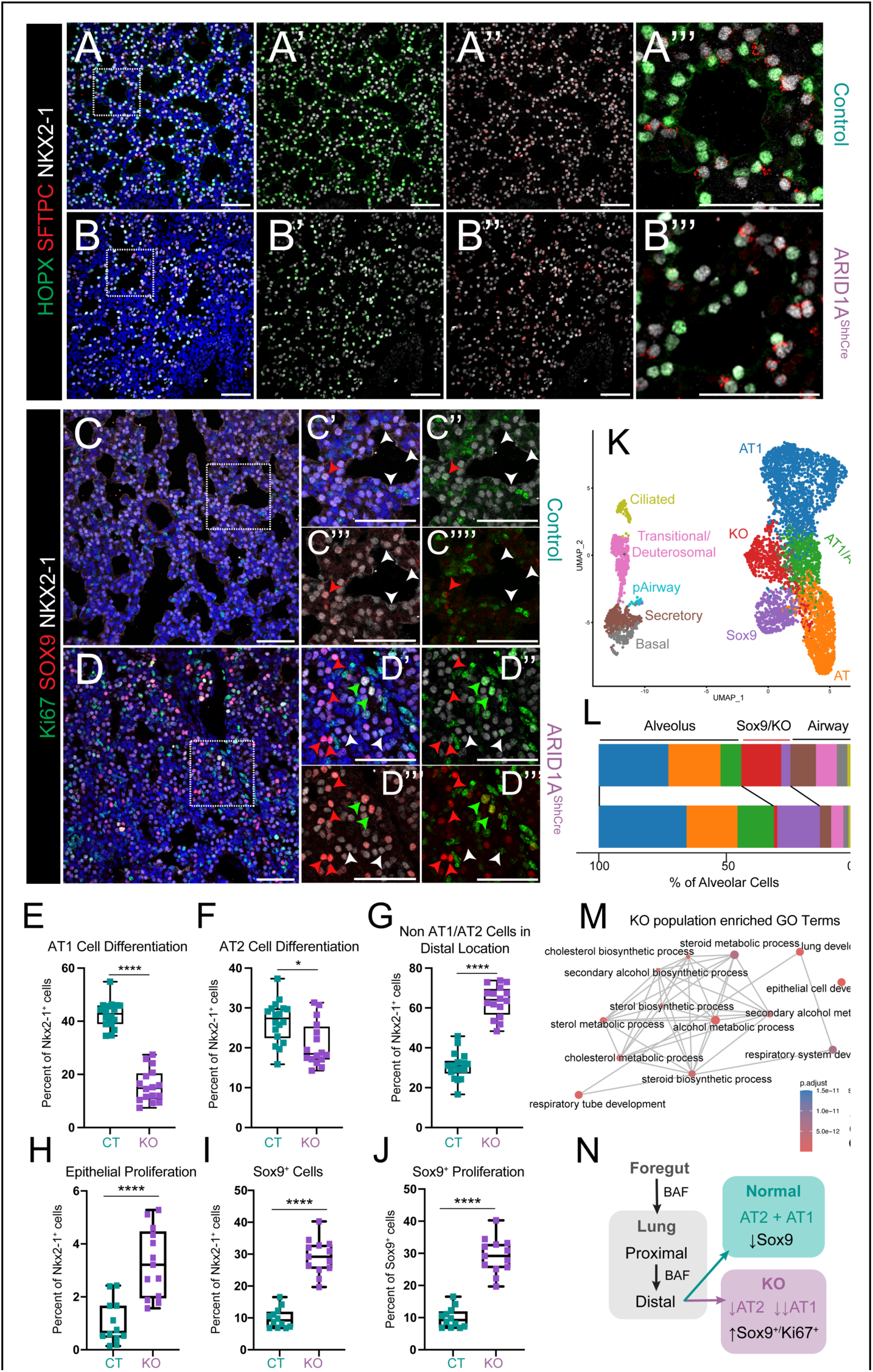
ARID1A and the BAF complex is required for distal epithelial cell specification in the embryonic lung. **(A-D)** Immunostaining for HOPX, SFTPC, and NKX2-1 marker **(A-B)** and NKX2-1, KI67 (proliferative marker) and SOX9 **(C-D)** in Control and ARID1A^ShhCre^ lungs at E18.5 **(E-J)** Quantification of cell states seen in A-D. **(K)** Epithelial clusters from scRNA-seq of Control and ARID1A^ShhCre^ lungs at E18.5 (**L)** Distribution of cell populations in K. (**M)** GO Biological Process Terms enriched in the KO cell population. **(N)** Model of BAF function in distal fate specification. Scale bars, 50 μm.

### Emergence of YAP/TAZ^Hi^ and Wnt^Hi^ cell population following deletion of ARID1A in the embryonic respiratory epithelium

Extensive prior literature supports the role of multiple signaling pathways in AT1 vs AT2 cell fate allocation in distal lung differentiation. Recent reports indicate that ARID1A directly interacts with YAP in the cytoplasm, and that loss of ARID1A protein expression leads to increased YAP nuclear localization *in vitro*^38^. Furthermore, active nuclear YAP is a hallmark of AT1 cells in both development and homeostasis in the lung^5,39^. We therefore reasoned that ARID1A loss may be associated with YAP dysregulation following the loss of epithelial ARID1A and BAF complex activity (Figure 6). Indeed, we observed that nuclear YAP and TAZ are typically present in AT1 cells in control embryos (Figure 6B,D), with over 80% of YAP^+^ or TAZ^+^ epithelial cells expressing HOPX in control lungs (Figure 6H,I). However in ARID1A^ShhCre^ mice, we noted a significant decrease in HOPX in YAP^+^ or TAZ^+^ cells (Figure 6C,E,H,I), with a corresponding increase in SOX9 in YAP^+^ or TAZ^+^ cells. Expression of a module of YAP/TAZ target genes from scRNAseq predicted YAP signaling response in the SOX9^+^ ARID1A^ShhCre^-specific epithelial cluster, whereas YAP response was present only in AT1 cells in control mice (Figure 6K). These data support the failure of AT1 differentiation in ARID1A^ShhCre^ mice despite increased YAP/TAZ pathway activity.

**Figure 6.**
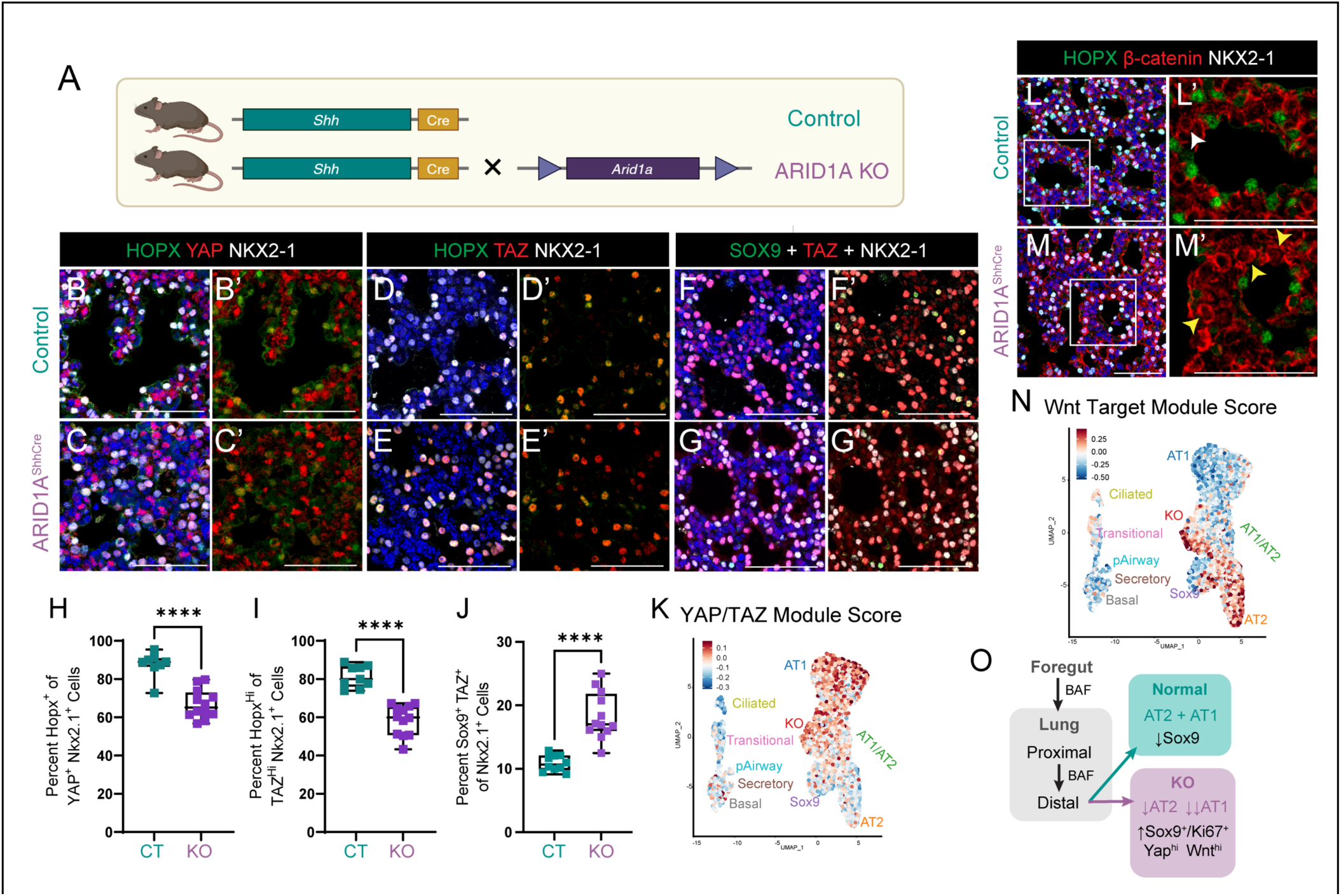
Persistence of YAP/TAZ^Hi^ and Wnt^Hi^ cell populations following deletion of ARID1A in the embryonic respiratory epithelium. **(A)** Genetic ablation of ARID1A in the embryonic respiratory epithelium. (**B-G)** Immunostaining of YAP, HOPX, NKX2-1, TAZ, and SOX9 in Control and ARID1A^ShhCre^ mice. **(H-J)**. Quantification of cell populations defined in B-G. **K**. YAP/TAZ target gene module score in scRNAseq. **(L-M)** Immunostaining of non-phosphorylated (active) β-catenin, HOPX, and NKX2-1. Yellow arrowheads show areas of increased nuclear β-catenin in ARID1A^ShhCre^ mice. **N**. Wnt target gene module score in scRNAseq. **(O)** Model of BAF function in distal fate signaling response. Scale bars, 50 μm.

WNT signaling is also required for AT1 vs AT2 differentiation, with high WNT signaling driving AT2 fate and WNT inhibition increasing AT1 fate^40^. Loss of ARID1A in other endodermal organs increases WNT signaling though β-catenin activation^41^. To assess WNT responsiveness in ARID1A^ShhCre^ mice, we evaluated nuclear localization of non-phosphorylated (active) β-catenin, a marker of active WNT response. Nuclear β-catenin was increased in the epithelium of ARID1A^ShhCre^ mice compared to control (Figure 6L-M, arrowheads), and expression of a set of direct WNT target genes was increased in the ARID1A^ShhCre^-specific KO epithelial cluster by scRNAseq. In contrast, these genes were highly expressed only in AT2 cells in control mice (Figure 6N). Notably, HOPX^+^ AT1 cells present in both control and ARID1A^ShhCre^ had no active nuclear β-catenin, demonstrating that cells which acquired AT1 cell identity in ARID1A^ShhCre^ mice properly downregulated WNT signaling. Taken together, these data support the hypothesis that aberrant simultaneous activation of the YAP/TAZ and WNT signaling pathways may prevent distal epithelial specification in ARID1A^ShhCre^ mice (Figure 6O).

### Failure of distal epithelial specification mispatterns of mesenchymal and endothelial compartments in the embryonic lung

Next, we evaluated the consequences of altered epithelial patterning on the other compartments of the developing lung. To evaluate changes in the mesenchymal compartment, we performed immunostaining at E18.5 for MEOX2 and ACTA2 in the developing saccular lung. In control lungs, we observed MEOX2 and ACTA2 immunostaining in distinct populations (Figure 7A). In ARID1A^ShhCre^ mice, there were more ACTA2^+^ cells and a markedly increased number of MEOX2^+^/ACTA2^+^ cells (Figure 7B). Concordantly, scRNA-seq analysis of mesenchymal cell populations at E18.5 indicated an increase in *Meox2*^+^/*Acta2*^+^ cells in the ARID1A^ShhCre^ mesenchyme (Figure 7C-D). Together, these data demonstrated non-cell autonomous failure of mesenchymal differentiation and suggested a myogenic preference in MEOX2^+^ the distal mesenchyme of ARID1A^ShhCre^ lungs.

**Figure 7.**
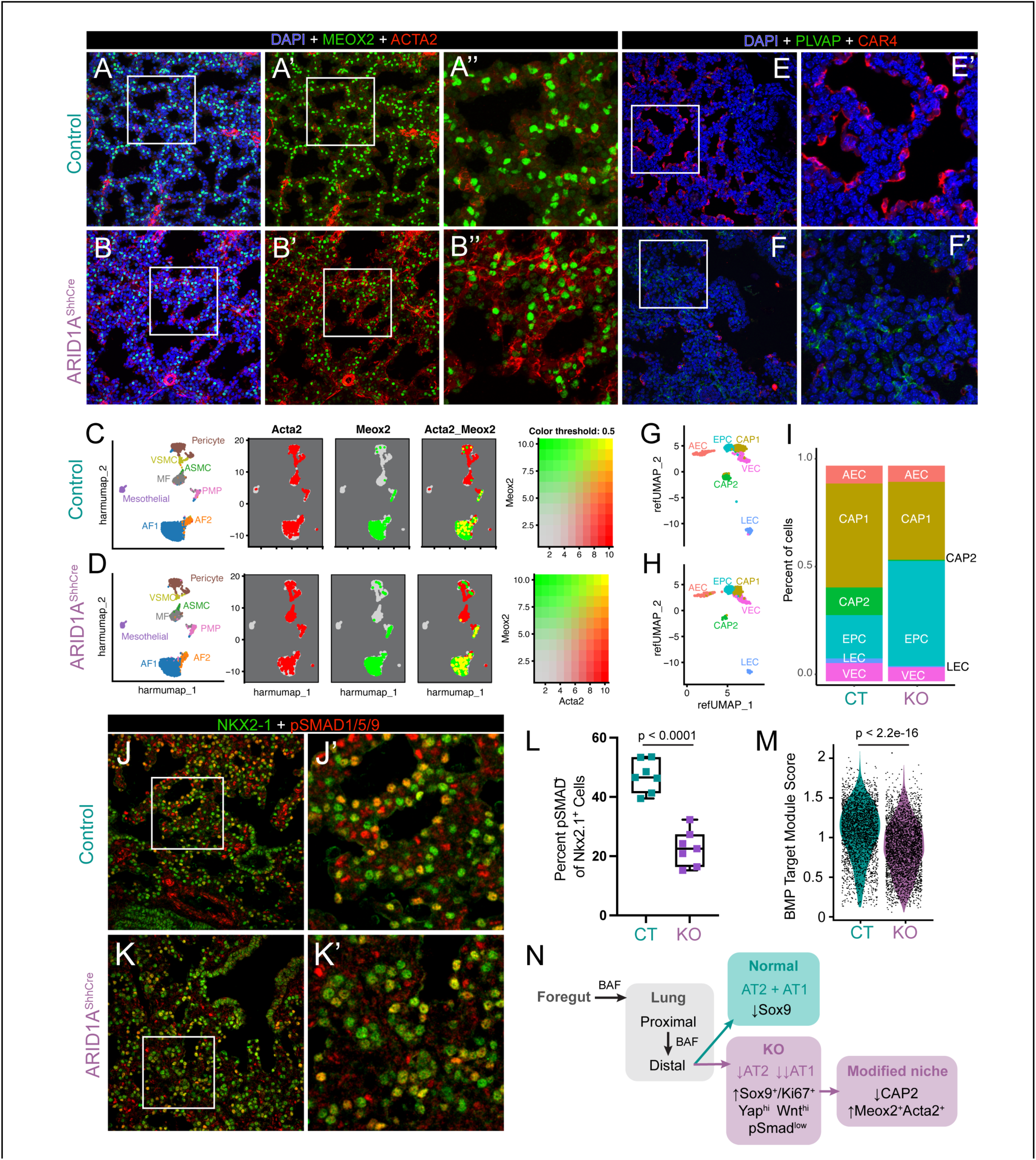
Failure of distal epithelial specification leads to failure of mesenchymal and endothelial patterning in the embryonic lung. **(A-B)** Immunostaining of mesenchymal cells (MEOX2, ACTA2) in control and ARID1A^ShhCre^ lungs at E18.5. **(C-D)** Single-cell RNA sequencing (scRNA-seq) of control and ARID1A^ShhCre^ lungs at E18.5 with mesenchymal clusters shown. Co-expression of *Meox2* and *Acta2* in scRNA-seq. **(E-F)** Immunostaining of capillary endothelial cells (CAR4, PLVAP) in control and ARID1A^ShhCre^ lungs at E18.5. **(G-H)** scRNA-seq with only endothelial clusters shown. (**I)** Proportion of cells in each endothelial cluster. **(J-K)** Immunostaining of phosphorylated (active) SMAD1/5/9 and lung epithelial lineage marker (NKX2-1). **(L)** Quantification of imaging in J-K. (**M)** BMP target gene module score from epithelial scRNA-seq. (**N)** Model of BAF function in distal fate development including modified niche. Scale bars, 50 μm.

We then evaluated the endothelial compartment by performing immunostaining for PLVAP to identify capillary 1 (CAP1) and CAR4 to identify capillary 2 (CAP2) cells of the developing endothelium. We observed a marked reduction in CAP2 cells and a corresponding increase in CAP1 and EPC cells in the ARID1A^ShhCre^ lung endothelium (Figure 7E-F), consistent with previous reports demonstrating the role of AT1 cells in providing signals critical to CAP2 differentiation^42^. scRNA-seq analysis of endothelial cell populations confirmed a large decrease in CAP2 cells, with near absence in ARID1A^ShhCre^ mice (Figure 7G-I). These data demonstrated critical defects in mesenchymal and endothelial cells as a consequence of impaired distal epithelial differentiation, creating a modified niche in the distal lung of ARID1A^ShhCre^ mice.

To further explore this modified niche consisting of increased MEOX2^+^/ACTA2^+^ cells and absent CAP2 cells, we evaluated the status of additional key signaling factors which impact distal epithelial development including FGF, TGF-β and BMP4 signaling. While scRNAseq module scoring showed no change in FGF and TGF-β signaling in ARID1A^ShhCre^ epithelium (data not shown), a module of direct BMP4 signaling targets was significantly decreased across all epithelial cells (Figure 7L). Consistent with this prediction, evaluation of BMP signaling response in epithelial cells by immunostaining for phosphorylated (p)SMAD1/5/9 demonstrated a decrease in pSMAD1/5/9^+^/Nkx2-1^+^ epithelial cells in ARID1A^ShhCre^ mice (Figure 7K). Taken together, these findings support the concept that changes in distal epithelial cell specification disrupt both cell differentiation and signaling networks in the distal lung niche of ARID1A^ShhCre^ mice (Figure 7M).

### BMP signaling rescues distal differentiation in embryonic lung organoids derived from

*ARID1A*^*ShhCre*^ *lungs*. As demonstrated in Figure 7, loss of epithelial ARID1A-associated BAF complex activity caused changes in YAP/TAZ, Wnt, and BMP4 signaling in the developing distal epithelium, with ARID1A^ShhCre^ mice harboring a unique YAP^hi^, WNT^hi^, pSMAD^low^ epithelial population. We hypothesized that BAF complex activity would be required for appropriate response to these signaling pathways, and failure of BAF complex activity was the primary reason ARID1A-null cells were unable to differentiate appropriately. To test this hypothesis, we modified a previously reported protocol for culturing adult lung alveolar organoids to grow embryonic murine lung organoids^43^. In brief, E18.5 EPCAM^+^ cells were sorted by magnetic-activated cell sorting (MACS) and seeded together with previously expanded WT E18.5 fibroblasts and Matrigel. Control lung organoids grown using this method showed robust colony formation (Figure 8A-B), and after 21 days in culture were of modest size (Figure 8C) with central cavitation and clear differentiation of RAGE^+^ AT1 cells and SFTPC^+^ AT2 cells (Figure 8D). Lung organoids grown from ARID1A^ShhCre^ epithelium cultured with WT mesenchyme exhibited decreased colony formation but increased size (Figure 8A-C). Whole-mount immunostaining for RAGE, SFTPC, and EdU showed a substantial increase in cell proliferation and near absence of AT1 or AT2 differentiation in ARID1A^ShhCre^ epithelium, recapitulating key characteristics of altered distal respiratory epithelium of ARID1A^ShhCre^ mice seen *in vivo* (Figure 8D-E). We therefore concluded that embryonic lung organoids provided a system to study the interaction of signaling pathway modulation and BAF complex activation in epithelial fate acquisition.

**Figure 8.**
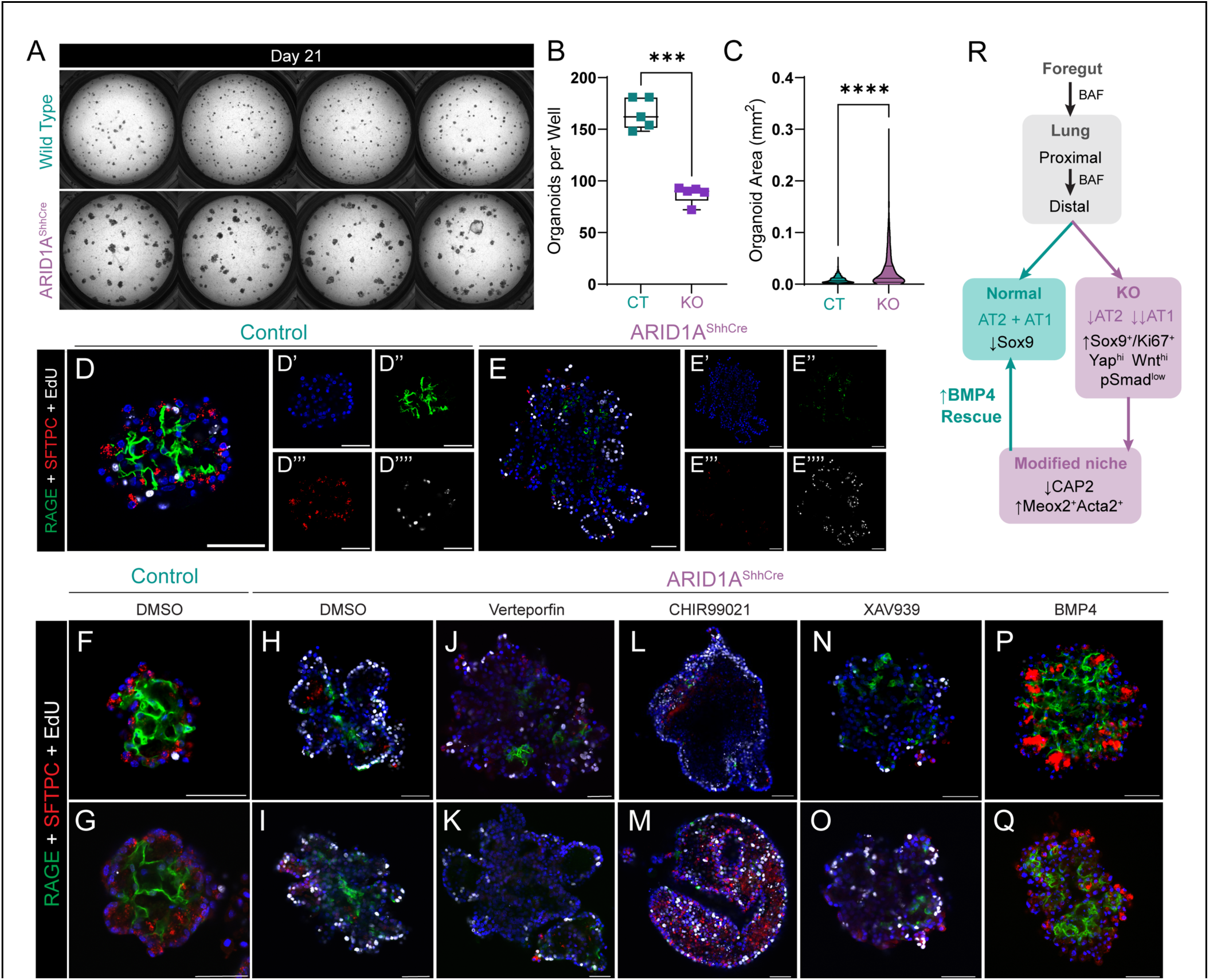
Exogenous BMP4 rescues distal epithelial differentiation in murine embryonic lung organoids. **A**. Brightfield imaging of embryonic murine lung organoids cultured from EPCAM^+^ cells isolated from control and *Arid1a*^fl/fl^; *Shh*^Cre^ lungs at E18.5. **B, C**. Quantification of number and size of organoids from imaging in A. **D, E**. Whole-mount immunostaining of RAGE, SFTPC, and EdU of representative Control and *Arid1a*^fl/fl^; *Shh*^Cre^ lung organoids. **F-Q**. Whole-mount immunostaining of RAGE, SFTPC, and EdU of representative Control and *Arid1a*^fl/fl^; *Shh*^Cre^ lungs organoids treated with DMSO, Verteporfin, CHIR99021, XAV939, and BMP4. **R**. Final model of BAF function in embryonic lung development. Scale bars, 50 μm.

We then tested whether modulation of YAP/TAZ, WNT, and BMP4 signaling could improve the differentiation defects of ARID1A^ShhCre^ knockout epithelium in organoid culture. We treated control and ARID1A^ShhCre^ embryonic lung organoids with DMSO, Verteporfin (a strong YAP inhibitor), CHIR99021 (an activator of canonical WNT signaling), XAV939 (an inhibitor of canonical WNT signaling), or recombinant BMP4 beginning on day 4 of organoid culture (Figure 8F-Q). Inhibition of YAP/TAZ signaling did not significantly alter AT1 or AT2 cell differentiation (Figure 8J-K). While activation of WNT signaling via CHIR99021 increased cell proliferation, it did not increase differentiation of AT2 cells (Figure 8L-M) and inhibition of WNT signaling did not increase AT1 cell differentiation (Figure 8N-O). In contrast, recombinant BMP4 treatment dramatically increased both AT1 cell and AT2 cell differentiation in ARID1A^ShhCre^ epithelium (Figure 8P-Q), with near complete rescue of organoid morphology and cellular quorum. Together, these data demonstrate that ARID1A and BAF complex activity is required for appropriate gene regulation downstream of YAP/TAZ and WNT signaling in distal lung epithelium. In contrast, BMP4 signaling can rescue impaired distal epithelial differentiation caused by the loss of epithelial ARID1A (Figure 8R), implying an alternative mechanism independent of or downstream of ARID1A-containing BAF complex function.

## Discussion

Cell fate acquisition and differentiation through the proximal-distal axis of the lung are critical processes required for respiratory system development. Chromatin modification changes genome structure to alter TF binding and regulate of transcriptional programs that drive fate acquisition. Here, we show that the canonical BAF complex, a core chromatin regulatory complex which facilitates H3K27 acetylation to increase chromatin accessibility, is required for lung endoderm differentiation and proximal-distal axis specification critical for normal lung formation. Combined knockout of epithelial ARID1A/1B to ablate BAF function prior to lung specification caused failure of lung bud formation, emphasizing the centrality of BAF complex function in lung development. Our finding that deletion of ARID1A has a more severe developmental phenotype than ARID1B deletion demonstrates both that ARID1A is the predominant subunit required in lung epithelial development and that ARID1A/B are partially redundant in lung endoderm differentiation since ARID1A-containing BAF complex is required for proximal-distal specification and appropriate alveolar fate acquisition in distal lung progenitors, and loss of proper epithelial specification drives altered patterning of lung mesoderm and endothelium. Finally, our data demonstrate that WNT and YAP/TAZ signaling require robust BAF complex activity to drive fate acquisition in the distal lung, and that BMP signaling can rescue both AT1 and AT2 differentiation, even in the absence of ARID1A.

The canonical BAF complex is only one of three known subtypes of mammalian SWI/SNF complexes. Prior data implicated SMARCA4 in the epithelial response to injury in adult lung and in the pathogenesis of lung cancer in both murine models and human NSCLC^44-47^. Like ARID1A/1B, SMARCA4 and SMARCA2 are mutually exclusive and partially redundant subunits, but are found in all three mammalian subtypes. Identification of the precise roles of each of these large and diverse complexes is a challenge, as even the delineation between the role of ARID1A-containing and ARID1B-containing BAF complexes during lung development is not entirely evident from this single study focused on genetic knockout as a primary assessment of gene activity. These complexes share several traits, such as their opposition to PRC2 complex activity and their essential role in nucleosome ejection^13,48^. A common limitation in studies of core regulatory pathways including BAF complex is that these pathways are used in most cells which in turn act pleiotropically on many aspects of cell state. Our data, however, imply more specificity for BAF complex activity in lung than would have been previously expected. Airway-specific knockout of ARID1A with *Sox2*^CreER^ does not disrupt proximal patterning, defining the key domain for BAF function as the distal lung epithelium after initial specification during lung organogenesis. These data suggest that studies focused on the cell- and tissue-specific roles of each subtype of BAF complex are may be fruitful to better define mechanisms of cell identity in the lung.

Present data add to a growing body of literature focused on the requirement for a balance of multiple chromatin regulators in lung epithelial development, regeneration, and disease. Deletion of EZH2, a core subunit of the PRC2 complex, during early lung developments causes expansion of basal cells which is reminiscent of phenotypes reported here in ARID1A^ShhCre^ animals^49^. BAF and PRC2 are well-known antagonists, with PRC2 catalyzing deposition of H3K27 trimethylation marks to inhibit transcription and compact chromatin^50^. It is intriguing to consider that loss of both complexes seems to prevent distal lung specification, supporting a common need for both appropriate activation and repression of key regulatory elements to properly define distal lung epithelial fate. Epithelial deletion of BRD4, an epigenetic reader that evicts nucleosomes from chromatin, also impairs distal epithelial differentiation, as noted by decreased AT2 cell numbers and a failure of maturation of AT1 cells^51^. BRD4 physically binds to and interacts with ARID1A to promote cell fate specification and differentiation in osteoclasts^52,53^; it is tempting to speculate that a similar interaction may underlie similar phenotypes in BRD4 and ARID1A knockouts in the lung epithelium. It was recently reported that CDK8/CDK19 kinase module phosphorylates ARID1A and other mammalian SWI/SNF complex components and deletion of these Mediator kinases caused a near-complete loss of ARID1A genomic occupancy^54^. In the same study, the Mediator subunit MED12 physically interacted with ARID1A in Co-IP assays in intestinal organoids. Together, these observations raise the possibility that ARID1A and the BAF complex is itself subject to another level of tightly controlled transcriptional and epigenetic regulation by the Mediator complex and other epigenetic modifiers such as BRD4.

One such epigenetic complex may be the histone methyltransferases PRDM3 and PRDM16. We recently reported that PRDM3/16 are required in the lung epithelium for AT2 cell lineage specification and differentiation during embryonic lung development^55^. Deletion of PRDM3/16 led to failure of AT2 differentiation with largely intact AT1 allocation, the inverse of the differentiation defect seen in ARID1A knockouts in the present study. Pathway analysis of differentially expressed AT1 cell associated genes predicted ARID1A and SMARCA4 as upstream regulators of the NKX2-1/PRDM3/PRDM16 centered transcriptional regulatory network in AT1 cells, which is consistent with the data presented here implicating the canonical BAF complex in AT1 differentiation. Together, these observations suggest that PRDM3/16 and the BAF complex may work in concert to balance the AT1 and AT2 cell differentiation to properly pattern the distal lung during the saccular stage of development, perhaps via direct binding to NKX2-1. This model is supported by the recent finding that loss of NKX2-1 in AT2 cells leads to acquisition of transcriptional and epigenetic features of AT1 fate, while the converse is true in AT1 cells^7^. This interpretation is also consistent with lineage tracing observations implying early separation of AT1 and AT2 lineages in distal lung development^56^. Given the proposed balance during embryonic development, we speculate that the BAF complex and the emerging network of coordinating partners may be common to alveologenesis, postnatal lung homeostasis, and lung regeneration following acute lung injury.

Chromatin regulators are generally thought to be ubiquitous and act similarly in all cell types. It is therefore provocative that BMP signaling mediated by SMADs does not require BAF activity to drive AT1 and AT2 differentiation in organoids, while WNT and YAP signaling are dependent on functional BAF in lung epithelium. One can speculate on multiple potential mechanisms underlying this pathway-related specificity. Perhaps WNT and YAP require “more” BAF activity than BMP signaling, allowing residual ARID1B-containing BAF to overcome ARID1A knockout for SMAD-dependent regulation. Alternatively, WNT and YAP could require ARID1-containing canonical BAF, while BMP-dependent SMAD requires alternative BAF complex isoforms or differential chromatin modifiers to drive gene expression in differentiation. In either case, our data highlight an unexpected specificity of signaling pathways mediated by BAF in the distal lung and imply that the tuning of cell fate in biological systems includes more complexity than a classical model of “core” chromatin regulators may imply. Future studies to understand these dynamics may therefore help chart a path for specificity in targeting chromatin regulators for translational and therapeutic purposes despite their apparent ubiquity.

## Acknowledgements

The authors would like to thank the Bio-Imaging and Analysis Facility, the Single Cell Genomics Facility, and the Genomic Sequencing Facility of the Cincinnati Children’s Research Foundation for technical support and access to key protocols.

## Author Contributions

Conceptualization – XS, DS, DTS, WJZ

Data acquisition – HL, AJ, SF, BZ, AS, ST, NB-A, KC, SH, SG

Data analysis – HL, AJ, SF, DS, XS, DTS, WJZ

Supervision – AZ, SB, JW, XS, DS, DTS, WJZ

Writing – original draft – HL, SF, WJZ

Writing – review and editing – all authors

## Data and materials availability

All single cell data is uploaded to the GEO database and has been uploaded to the LungMAP ingest server for archiving on lungmap.net. All other data is available by request to the corresponding author.

## Materials and Methods

### Ethical compliance and animals

All animal studies were conducted under the guidance and supervision of the Cincinnati Children’s Hospital Medical Center (CCHMC) Institutional Animal care and Use Committee (IACUC) in accordance with CCHMC regulatory and biosafety protocols. All experimental mice were bred and maintained on a mixed C57BL/6, CD-1 background. Mouse lines used include: *Arid1a*^*tm1*.*1Zhwa*^/J (JAX strain 027717), *Arid1b*^*em2Hzhu*^/J (JAX strain 032061), *Shh*^*tm1(EGFP/cre)Cjt*^/J (JAX strain 005622), *Sox2*^*tm1(cre/ERT2)Hoch*^/J (JAX strain 017593), and CD-1 (Charles River strain 022). For *Shh*^*Cre*^ breeding, Arid1a and Arid1b genes were conditionally deleted individually or in combination by crossing *Arid1a*^*flox/+*^*Shh*^*Cre*^, *Arid1b*^*flox/+*^*Shh*^*Cre*^ *Arid1a*^*flox/+*^*Arid1b*^*flox/+*^*Shh*^*Cre*^ males corresponding *Arid1a*^*flox/flox*^, *Arid1b*^*flox/flox*^ and *Arid1a*^*flox/flox*^*Arid1b*^*flox/flox*^ females. Embryos were harvested at E11.5, E12.5, E14.5, E16.5 and E18.5 respectively. For the Sox2-CreER breeding, *Arid1a*^*flox/flox*^*Sox2*^*CreERT2*^ females were crossed with *Arid1a*^*flox/flox*^ male mice. Animals were mated overnight, and the presence of a vaginal plug the next morning was defined as E0.5. To induce and with CreER activity, *Sox2*^*CreERT2+*^ pregnant dams were given one dose of 10 mg tamoxifen dissolved in peanut oil (100 μL of 100 mg/mL) at E12.5 via oral gavage and embryos were harvested at E18.5. Controls in all cases included littermates lacking Shh-Cre or Sox2-CreER expression.

### Mouse lung harvest

Pregnant dams were euthanized via CO_2_ followed by cervical dislocation and thoracotomy. Embryos isolated at E16.5 or later were euthanized by decapitation. Lungs were isolated with extrapulmonary tissue removed.

### Histology and immunofluorescence

Embryonic lungs were harvested and fixed in 4% paraformaldehyde in PBS at 4 °C overnight. Tissues were washed with PBS and dehydrated through and ethanol gradient and processed overnight using a standardized automated processing protocol (Thermo Scientific, Excelsior ES). The samples were embedded in paraffin and sectioned at 5 μm thickness. Paraffin sections were stained with Hematoxylin and Eosin using standard protocols and imaged on a Nikon Eclipse Ti2 microscope. Immunofluorescence staining on paraffin sections was performed using previously described protocols^43,57^. The primary antibodies used include: Anti-Arid1a (Rabbit, Abcam, ab182560 – 1:100), Anti-Arid1b (Mouse IgG2b, Abcam, ab57461 – 1:200), Anti-TTF1/Nkx2.1 (Guinea Pig, Seven Hills Bioreagents, G237 - 1:100), Anti-Sox2 (Mouse IgG1, Santa Cruz, sc-365823 – 1:100), Anti-Tub1A1 (Mouse IgG2b, Millipore, T7451 - 1:1000), Anti-Scgb1a1/CCSP (Rabbit, Seven Hills, WRAB-3950 - 1:500), Anti-P63 (Mouse, Biocompare, CM163A - 1:100), Anti-Krt5 (Rabbit, Biolegend, 905903 - 1:100), Anti-HOPX (Mouse IgG1, Santa Cruz, sc-398703 - 1:100), Anti-SFTPC (Guinea Pig, Seven Hills Bioreagents, G992 - 1:500), Anti-Sox9 (Rabbit, EMD Millipore, AB5535 - 1:100), Anti-Ki67 (Mouse IgG1, BD Biosciences, 556003 - 1:100), Anti-Car4 (Goat, R&D Biosystems, AF2414 - 1:200), Anti-PLVAP (Rat, Bio-Rad, MCA2539GAA, MECA-32 - 1:100), Anti-ACTA2 (Mouse, Sigma, A5228 - 1:2000), Anti-MeOX2 (Rabbit, Novus Biologicals, NBP2-30647 - 1:100). The corresponding secondary antibodies (1:200) used include: Goat anti-Guinea Pig IgG Alexa Fluor 647 (Invitrogen, A21450), Goat anti-Mouse IgG1 Alexa Fluor 488 (Invitrogen, A21121), Goat anti-Rabbit IgG Alexa Fluor 568 (Life Technologies, A11036), Goat anti-Mouse IgG2b Alexa Fluor 568 (Invitrogen, A21144), Goat anti-rat Alexa Fluor 647 (Invitrogen, A21247), Goat anti-Goat IgG Alexa Fluor 568 (Invitrogen, A11057) and Goat anti-Mouse IgG Alexa Fluor 568 (Invitrogen, A11001). Following secondary antibody application, the sections were quenched to remove autofluorescence using Vector TrueVIEW Autofluorescence Quenching kit (Vector labs, Cat# SP-8400), stained with DAPI (1:2000) and mounted using Prolong Gold antifade mounting medium (Invitrogen, P36930). Immunofluorescence for Yap (Rabbit, Cell Signaling Technology, 4912 - 1:100), Taz (Rabbit, Cell Signaling Technology, 83669 - 1:200), p-Smad1/5/9 (Rabbit, Cell Signaling Technology, 13820T, 1:100) and non-phospho-b-catenin (Rabbit, Cell Signaling Technology, 8814T - 1:500) was performed using ImmPRESS® HRP Horse Anti-Rabbit IgG Polymer Detection Kit (Vector Labs, MP-7401-50) which was followed Tyramide signal amplification (TSA Plus Cyanine 3.5, Akoya Biosciences, NEL763001KT) with an 8min exposure time. Following the application of TSA fluorophores, sections were stained with DAPI (Invitrogen, D1306; 1:1000) and mounted using Prolong Gold antifade mounting medium (Invitrogen, P36930).

For AT1, AT2 cell quantification, four 40X Z-stack confocal images were taken at the distal margins for each control and knockout (n=3). All Z-stack images were maximum intensity projected in FIJI software and were used to manually score AT1/ AT2 cells. To reduce the autofluorescence background signal created by red blood cells, we used FIJI to create a new channel which contained only auto fluorescent blood cells by subtracting the FITC channel from the Cy5 channel and then using the newly created FITC-Cy5 channel to subtract out the blood cells from the FITC and TRITC channels. All epithelial cells (DAPI^+^/NKX2.1^+^) were first manually counted and then the AT1 (DAPI^+^/NKX2.1^+^/HOPX^+^) cell subset and AT2 (DAPI^+^/NKX2.1^+^/SFTPC^+^) cell subsets were manually counted. Similarly, manual counts were done for the following immunofluorescent signals: SOX2, YAP, TAZ, SOX9, Ki67 and p-SMAD1/5/9.

### Whole-mount immunofluorescence

Tracheal lung tissues isolated at E11.5 and E12.5 were subject to whole mount immunofluorescence as previously described^58^. Embryonic tissue was fixed in 4% PFA overnight and then stored in 100% Methanol (MeOH) at -20°C. For staining, wholemounts were permeabilized in Dent’s Bleach (4:1:1 MeOH: DMSO: 30% H_2_O_2_) for 2 hours, then taken from 100% MeOH to 100% PBS through a series of washes. Following washes, wholemounts were blocked in a 2% BSA (w/v) blocking solution for two hours and then incubated, overnight, at 4 °C in primary antibody diluted accordingly in the blocking solution. After five one-hour washes in PBS, wholemounts were incubated with a secondary antibody at a dilution of 1:500 overnight at 4 °C. Samples were then washed three times in 1X PBS, transferred to 100% methanol, through a series of washes in dilutions of methanol, and cleared in benzyl-alcohol benzyl-benzoate (Murray’s Clear). Images of wholemounts were obtained using confocal microscopy (Nikon A1R). Imaris imaging software was used to convert z-stack image slices obtained using confocal microscopy to 3D renderings of wholemount samples.

### Mouse lung single cell suspension preparation and sequencing

E10.5 lungs were dissected, and extrapulmonary tissue was removed from the embryos. For each sample, three lungs of the same genotype were finely chopped and placed into Eppendorf tubes. The samples were digested in 1 mL of 10X TryPLE solution (Gibco, A1217701) at 37 °C for 10minutes with intermittent pipetting to dissociate cells. Each sample was then passed through a 35 μm filter into a falcon tube (Fisher Scientific, 08-771-23) and placed on ice. Cells were pelleted by centrifugation at 300xg for 5 minutes at 4 °C and the supernatant was discarded. The pellets were resuspended in 100 μL of HBSS+FBS (equal volumes) and transferred to clean eppendorf tubes. Cells were counted using a hemocytometer in the presence of trypan blue, and the concentration was adjusted to 1000 cells/μL for downstream processing for scRNAseq.

E18.5 lungs were dissected, and extrapulmonary tissue was removed from the embryos. For each sample, two lungs of the same genotype were minced together using a razor blade to create a fine paste and transferred to 10 mL conical tubes. Each sample was digested for 30 min at 37 °C in 3ml of Collagenase/Dispase/DNase I enzyme mix (6.3 mg of Collagenase Type I [Invitrogen, 17100-017], 300 μL Dispase [Corning, 354235], 30 μL DNaseI [BD Bioscience, 354235, 10,000 U/mL] added upto 3 mL of Dulbecco’s phosphate buffered saline [Thermo Fisher, 14190144]) with intermittent vortexing and pipetting. Samples were then passed through a 40 μm filter (Greiner Bio-One, 542040) and centrifuged at 500×*g* for 5 min at 4 °C and the supernatant discarded. The cell pellets were resuspended in 1 mLof RBC lysis buffer (ACK lysis buffer, Gibco, A1049201) and placed on ice for 3 min. After RBC lysis, the samples were diluted with 3 mL of ice-cold PBS and centrifuged at 500×*g* for 5 min at 4 °C. Following removal of the supernatant, cells were resuspended in HBSS+FBS (equal volumes), counted, and resuspended at a final concentration of 1000 cells/μL (50,000 cells total in 50 μL) for downstream processing for scRNAseq.

For each single cell preparation described above, a maximum of 16,000 cells or nuclei were loaded into a channel of a 10x Genomics Chromium system by the Cincinnati Children’s Hospital Medical Center Single Cell Sequencing Core. Libraries for RNA (3v3) were generated following the manufacturer’s protocol. Sequencing was performed by the Cincinnati Children’s Hospital DNA Sequencing Core using Illumina reagents.

### Embryonic lung organoids

Raw Sequencing data was aligned to the mouse reference genome mm10 with CellRanger 4.0 to generate expression count matrix files. Output files were loaded into Seurat 5.0 using and processed using *SCTransform*. Cells with less than 2000 or more than 8000 features were filtered and cells were clustered using the standard Seurat workflow. Putative doublets were identified and removed using DoubletFinder^59^, and libraries from individual time points and treatments were integrated using *SelectIntegrationFeatures* and *IntegrateData* commands in Seurat. Following integration, cells were re-clustered, UMAP project generated, and samples identified based on expression similarity cell states defined by LungMAP. Module scoring was performed using *AddModuleScore* function in Seurat for gene sets indicated in the figures. Targets for WNT signaling module included Ccnd1, Myc, Axin2, Jun, Id2, Lgr5, Lrp5, and Fgfr2. Targets for YAP included Ckd1, Myc, Sox2, Axl, Birc5, Bcl2, Ccn1, Ccn2, Snai1, Snai2. Targets for BMP signaling included Id1, Id2, Id3, Nanog, Vegfa, Hand1, Emilin2, Grem12, Nog, Fst, Smad6, and Smad7.

### Embryonic lung organoids

Embryonic lung organoids were generated in a similar manner to adult AEP-derived organoids as previously described with modifications as follows^43^. Single cell suspensions were generated from lungs harvested from Control and ARID1A^ShhCre^ E18.5 embryos as previously described. EPCAM^+^/CD45^-^/CD16/32^-^/CD90.2^-^/CD31^-^/LNGFR^-^ cells were isolated by magnetic activated cell sorting (MACS) as follows. The single cell suspension was resuspended in 5 mL MACS Buffer (autoMACS Rinsing Solution [Miltenyi Biotec, 130-091-222] with MACS BSA Stock Solution [Miltenyi Biotec, 130-091-376]) and passed through a 40 μm filter. Cells were centrifuged and resuspended in 500 μL MACS Buffer containing biotin anti-mouse CD45 antibody (BioLegend, 103104 – 1:100), biotin anti-mouse CD16/CD32 (BD Biosciences, 553143 – 1:100), biotin anti-mouse TER-119/erythroid cell antibody (BioLegend, 116204 – 1:200), biotin anti-mouse CD90.2 antibody, 105304 – 1:200), biotin anti-mouse CD31 antibody (BioLegend, 102504 – 1:200), and biotin mouse CD271 (LNGFR) antibody (Miltenyi Biotec, 130-110-110 – 1:250) for 30 min at 4 °C. Following centrifugation, cells were resuspended in 20 μL of anti-biotin microbeads (Miltenyi Biotec, 130-090-485) for 15 min at 4 °C followed by negative selection for CD45^-^/CD16/32^-^/CD90.2^-^/CD31^-^/LNGFR^-^ cells using a QuadroMACS separator (Miltenyi Biotec 130-090-976) and LS columns (Miltenyi Biotec 130-042-401). The resulting single cell suspension was then blocked with FcR for 10 min at 4 °C followed by incubation with EpCAM (CD326) microbeads (Miltenyi Biotec 130-105-958) for 15 min at 4 °C. A positive selection for EpCAM^+^ cells was performed and the resulting single cell suspension was resuspended in ‘spiked’ SAGM as previously described. Embryos for embryonic fibroblast stocks were isolated from wild-type pregnant CD-1 dams at E18.5. These stocks were generated, expanded, and thawed similarly to adult fibroblast stocks as previously reported. E18.5 EPCAM^+^/CD45^-^/CD16/32^-^/CD90.2^-^/CD31^-^/LNGFR^-^ cells and E18.5 fibroblasts were seeded at 5000 and 50000 total cells, respectively, with Corning Matrigel GFR Membrane Matrix (Corning, 356231) and maintained as previously described. Organoids were supplemented in the ‘spiked’ SAGM on day 4 of culture and onwards with Verteporfin (Tocris, 5305), CHIR99021 (Tocris, 4423), XAV939 (Tocris, 3748), and recombinant human BMP4 (R&D Systems, 314-BP).

### Organoid plate imaging and quantification

Whole-well imaging and quantification was performed as described previously^43^.

### Isolation of organoids for whole-mount immunohistochemistry

Previously reported protocols for whole-mount immunostaining of organoids were followed^43^. Primary antibodies used included anti-mouse/rat RAGE antibody (R&D Systems, MAB1179 – 1:100), anti-guinea pig SFTPC (Seven Hills Bioreagents, G992 - 1:100), and anti-guinea pig TTF1/Nkx2.1 (Seven Hills Bioreagents, G237 - 1:100) along with secondary antibodies diluted to 1:200 used for immunostaining of tissue sections described above. For whole-mount EdU staining, the Click-iT EdU Cell Proflieration Kit for Imaging, Alexa Fluor 647 dye (Invitrogen, C10340) was used in conjunction with the protocol for whole-mount immunohistochemistry as previously described^43^. Briefly, the organoid medium was replaced with ‘spiked’ SAGM with 10 μM EdU. The EdU signal was detected following the protocol from the commercial kit.

### Rapid immunoprecipitation mass spectrometry of endogenous proteins (RIME)

Mouse distal lung epithelial cell line (MLE-15) were cultured in HITES medium (RPMI 1640 (Gibco, 11875-093) supplemented with 5 μg/mL Insulin, Transferrin, Sodium Selenite (Sigma, I1884), 5μg/ml bovine apo-Transferrin (Sigma, T-1428), 10 nM hydrocortisone (Sigma, H0888), 10 nM *β*-estradiol (Sigma, E2758), 100 U/ml Penicillin / 100 μg/mL streptomycin (Sigma, P0781), 2mM glutamine (Sigma, G7513), 10 mM HEPES (Sigma, H0887) 2% Fetal bovine serum (FBS)). MLE-15 cells were grown in a monolayer to approximately 70% confluency and 50 million cells in duplicate were used for RIME submission using a previously described protocol^60^. For each sample, 50 million MLE15 cells were fixed in 1 mL Formaldehyde solution (11% Methanol free formaldehyde, 0.1 M NaCl, 1 mM EDTA, 50 mM HEPES in 10 mL of distilled water) for 8mins at room temperature. 2.5

M glycine was added to the cell media to stop the fixation and incubated for 5 min at room temperature. The cells were collected by scraping and transferring to a 15 mL conical tube and placed on ice. The cells were pelleted by centrifugation at 800 x *g* for 10mins at 4 °C, supernatant was discarded, and the cell pellets were resuspended in 10 mL chilled 1X PBS - 0.5% Igepal CA-630 (Sigma, I-8896) and 100 μL of 1 mM phenylmethanesulfonyl fluoride (PMSF) (Sigma #P-7626). The cells were then pelleted, the supernatant discarded, and the cell pellets were flash-frozen in liquid nitrogen and submitted to Active Motif for RIME analysis. Anti-Arid1A (Abcam, ab182560) was used for the RIME assay.

## References

1. Bellusci, S., Grindley, J., Emoto, H., Itoh, N., and Hogan, B.L.M. (1997). Fibroblast Growth Factor 10 (FGF10) and branching morphogenesis in the embryonic mouse lung. Development 124, 4867–4878. 10.1242/dev.124.23.4867.

2. Weaver, M., Yingling, J.M., Dunn, N.R., Bellusci, S., and Hogan, B.L.M. (1999). Bmp signaling regulates proximal-distal differentiation of endoderm in mouse lung development. Development 126, 4005–4015. 10.1242/dev.126.18.4005.

3. Shu, W., Guttentag, S., Wang, Z., Andl, T., Ballard, P., Lu, M.M., Piccolo, S., Birchmeier, W., Whitsett, J.A., Millar, S.E., and Morrisey, E.E. (2005). Wnt/β-catenin signaling acts upstream of N-myc, BMP4, and FGF signaling to regulate proximal–distal patterning in the lung. Developmental Biology 283, 226–239. 10.1016/j.ydbio.2005.04.014.

4. Lange, A.W., Sridharan, A., Xu, Y., Stripp, B.R., Perl, A.-K., and Whitsett, J.A. (2015). Hippo/Yap signaling controls epithelial progenitor cell proliferation and differentiation in the embryonic and adult lung. Journal of Molecular Cell Biology 7, 35–47. 10.1093/jmcb/mju046.

5. Nantie, L.B., Young, R.E., Paltzer, W.G., Zhang, Y., Johnson, R.L., Verheyden, J.M., and Sun, X. (2018). Lats1/2 inactivation reveals Hippo function in alveolar type I cell differentiation during lung transition to air breathing. Development 145, dev163105. 10.1242/dev.163105.

6. Volckaert, T., Yuan, T., Yuan, J., Boateng, E., Hopkins, S., Zhang, J.-S., Thannickal, V.J., Fässler, R., and De Langhe, S.P. (2019). Hippo signaling promotes lung epithelial lineage commitment by curbing Fgf10 and β-catenin signaling. Development 146, dev166454. 10.1242/dev.166454.

7. Little, D.R., Lynch, A.M., Yan, Y., Akiyama, H., Kimura, S., and Chen, J. (2021). Differential chromatin binding of the lung lineage transcription factor NKX2-1 resolves opposing murine alveolar cell fates in vivo. Nature Communications 12, 2509. 10.1038/s41467-021-22817-6.

8. Stuart, W.D., Fink-Baldauf, I.M., Tomoshige, K., Guo, M., and Maeda, Y. (2021). CRISPRi-mediated functional analysis of NKX2-1-binding sites in the lung. Communications Biology 4, 1–14. 10.1038/s42003-021-02083-4.

9. Rockich, B.E., Hrycaj, S.M., Shih, H.P., Nagy, M.S., Ferguson, M.A.H., Kopp, J.L., Sander, M., Wellik, D.M., and Spence, J.R. (2013). Sox9 plays multiple roles in the lung epithelium during branching morphogenesis. Proceedings of the National Academy of Sciences 110, E4456–E4464. 10.1073/pnas.1311847110.

10. Danopoulos, S., Alonso, I., Thornton, M.E., Grubbs, B.H., Bellusci, S., Warburton, D., and Al Alam, D. (2018). Human lung branching morphogenesis is orchestrated by the spatiotemporal distribution of ACTA2, SOX2, and SOX9. American Journal of Physiology-Lung Cellular and Molecular Physiology 314, L144–L149. 10.1152/ajplung.00379.2017.

11. Rawlins, E.L., Clark, C.P., Xue, Y., and Hogan, B.L.M. (2009). The Id2+ distal tip lung epithelium contains individual multipotent embryonic progenitor cells. Development 136, 3741–3745. 10.1242/dev.037317.

12. Zhang, Z., Newton, K., Kummerfeld, S.K., Webster, J., Kirkpatrick, D.S., Phu, L., Eastham-Anderson, J., Liu, J., Lee, W.P., Wu, J., et al. (2017). Transcription factor Etv5 is essential for the maintenance of alveolar type II cells. Proceedings of the National Academy of Sciences 114, 3903–3908. 10.1073/pnas.1621177114.

13. Clapier, C.R., Iwasa, J., Cairns, B.R., and Peterson, C.L. (2017). Mechanisms of action and regulation of ATP-dependent chromatin-remodelling complexes. Nature Reviews Molecular Cell Biology 18, 407–422. 10.1038/nrm.2017.26.

14. Alver, B.H., Kim, K.H., Lu, P., Wang, X., Manchester, H.E., Wang, W., Haswell, J.R., Park, P.J., and Roberts, C.W.M. (2017). The SWI/SNF chromatin remodelling complex is required for maintenance of lineage specific enhancers. Nature Communications 8, 14648. 10.1038/ncomms14648.

15. Mashtalir, N., D’Avino, A.R., Michel, B.C., Luo, J., Pan, J., Otto, J.E., Zullow, H.J., McKenzie, Z.M., Kubiak, R.L., St. Pierre, R., et al. (2018). Modular Organization and Assembly of SWI/SNF Family Chromatin Remodeling Complexes. Cell 175, 1272-1288.e1220. 10.1016/j.cell.2018.09.032.

16. He, S., Wu, Z., Tian, Y., Yu, Z., Yu, J., Wang, X., Li, J., Liu, B., and Xu, Y. (2020). Structure of nucleosome-bound human BAF complex. Science 367, 875–881. 10.1126/science.aaz9761.

17. Mashtalir, N., Suzuki, H., Farrell, D.P., Sankar, A., Luo, J., Filipovski, M., D’Avino, A.R., Pierre, R.S., Valencia, A.M., Onikubo, T., et al. (2020). A Structural Model of the Endogenous Human BAF Complex Informs Disease Mechanisms. Cell 183, 802-817.e824. 10.1016/j.cell.2020.09.051.

18. Gao, X., Tate, P., Hu, P., Tjian, R., Skarnes, W.C., and Wang, Z. (2008). ES cell pluripotency and germ-layer formation require the SWI/SNF chromatin remodeling component BAF250a. Proceedings of the National Academy of Sciences 105, 6656–6661. 10.1073/pnas.0801802105.

19. Hota, S.K., and Bruneau, B.G. (2016). ATP-dependent chromatin remodeling during mammalian development. Development 143, 2882–2897. 10.1242/dev.128892.

20. Liu, X., Dai, S.-K., Liu, P.-P., and Liu, C.-M. (2021). Arid1a regulates neural stem/progenitor cell proliferation and differentiation during cortical development. Cell Proliferation 54, e13124. 10.1111/cpr.13124.

21. Barnada, S.M., Gracia, A.G.d., Morenilla-Palao, C., López-Cascales, M.T., Scopa, C., Waltrich, F.J., Mikkers, H.M.M., Cicardi, M.E., Karlin, J., Trotti, D., et al. (2024). ARID1A-BAF coordinates ZIC2 genomic occupancy for epithelial to mesenchymal transition in cranial neural crest lineage commitment. 10.1101/2024.04.03.587869.

22. Han, L., Madan, V., Mayakonda, A., Dakle, P., Woon, T.W., Shyamsunder, P., Nordin, H.B.M., Cao, Z., Sundaresan, J., Lei, I., et al. (2019). Chromatin remodeling mediated by ARID1A is indispensable for normal hematopoiesis in mice. Leukemia 33, 2291–2305. 10.1038/s41375-019-0438-4.

23. Lei, I., Gao, X., Sham, M.H., and Wang, Z. (2012). SWI/SNF Protein Component BAF250a Regulates Cardiac Progenitor Cell Differentiation by Modulating Chromatin Accessibility during Second Heart Field Development*. Journal of Biological Chemistry 287, 24255–24262. 10.1074/jbc.M112.365080.

24. Wang, Z., Chen, K., Jia, Y., Chuang, J.-C., Sun, X., Lin, Y.-H., Celen, C., Li, L., Huang, F., Liu, X., et al. (2020). Dual ARID1A/ARID1B loss leads to rapid carcinogenesis and disruptive redistribution of BAF complexes. Nat Cancer 1, 909–922. 10.1038/s43018-020-00109-0.

25. Wang, W., Friedland, S.C., Guo, B., O’Dell, M.R., Alexander, W.B., Whitney-Miller, C.L., Agostini-Vulaj, D., Huber, A.R., Myers, J.R., Ashton, J.M., et al. (2019). ARID1A, a SWI/SNF subunit, is critical to acinar cell homeostasis and regeneration and is a barrier to transformation and epithelial-mesenchymal transition in the pancreas. Gut 68, 1245–1258. 10.1136/gutjnl-2017-315541.

26. Hiramatsu, Y., Fukuda, A., Ogawa, S., Goto, N., Ikuta, K., Tsuda, M., Matsumoto, Y., Kimura, Y., Yoshioka, T., Takada, Y., et al. (2019). Arid1a is essential for intestinal stem cells through Sox9 regulation. Proceedings of the National Academy of Sciences 116, 1704–1713. 10.1073/pnas.1804858116.

27. Cenik, B.K., and Shilatifard, A. (2021). COMPASS and SWI/SNF complexes in development and disease. Nature Reviews Genetics 22, 38–58. 10.1038/s41576-020-0278-0.

28. Centore, R.C., Sandoval, G.J., Soares, L.M.M., Kadoch, C., and Chan, H.M. (2020). Mammalian SWI/SNF Chromatin Remodeling Complexes: Emerging Mechanisms and Therapeutic Strategies. Trends in Genetics 36, 936–950. 10.1016/j.tig.2020.07.011.

29. Mittal, P., and Roberts, C.W.M. (2020). The SWI/SNF complex in cancer — biology, biomarkers and therapy. Nature Reviews Clinical Oncology 17, 435–448. 10.1038/s41571-020-0357-3.

30. Naito, T., Udagawa, H., Umemura, S., Sakai, T., Zenke, Y., Kirita, K., Matsumoto, S., Yoh, K., Niho, S., Tsuboi, M., et al. (2019). Non-small cell lung cancer with loss of expression of the SWI/SNF complex is associated with aggressive clinicopathological features, PD-L1-positive status, and high tumor mutation burden. Lung Cancer 138, 35–42. 10.1016/j.lungcan.2019.10.009.

31. Santen, G.W.E., Aten, E., Sun, Y., Almomani, R., Gilissen, C., Nielsen, M., Kant, S.G., Snoeck, I.N., Peeters, E.A.J., Hilhorst-Hofstee, Y., et al. (2012). Mutations in SWI/SNF chromatin remodeling complex gene ARID1B cause Coffin-Siris syndrome. Nature Genetics 44, 379–380. 10.1038/ng.2217.

32. Tsurusaki, Y., Okamoto, N., Ohashi, H., Kosho, T., Imai, Y., Hibi-Ko, Y., Kaname, T., Naritomi, K., Kawame, H., Wakui, K., et al. (2012). Mutations affecting components of the SWI/SNF complex cause Coffin-Siris syndrome. Nature Genetics 44, 376–378. 10.1038/ng.2219.

33. Qiao, L., Xu, L., Yu, L., Wynn, J., Hernan, R., Zhou, X., Farkouh-Karoleski, C., Krishnan, U.S., Khlevner, J., De, A., et al. (2021). Rare and de novo variants in 827 congenital diaphragmatic hernia probands implicate LONP1 as candidate risk gene. The American Journal of Human Genetics 108, 1964–1980. 10.1016/j.ajhg.2021.08.011.

34. Gofin, Y., Zhao, X., Gerard, A., Scaglia, F., Wangler, M.F., Schrier Vergano, S.A., and Scott, D.A. (2022). Evidence for an association between Coffin-Siris syndrome and congenital diaphragmatic hernia. American Journal of Medical Genetics Part A 188, 2718–2723. 10.1002/ajmg.a.62889.

35. Khattar, D., Fernandes, S., Snowball, J., Guo, M., Gillen, M.C., Jain, S.S., Sinner, D., Zacharias, W., and Swarr, D.T. (2022). PI3K signaling specifies proximal-distal fate by driving a developmental gene regulatory network in SOX9+ mouse lung progenitors. eLife 11, e67954. 10.7554/eLife.67954.

36. Duren, Z., Chen, X., Jiang, R., Wang, Y., and Wong, W.H. (2017). Modeling gene regulation from paired expression and chromatin accessibility data. Proceedings of the National Academy of Sciences 114, E4914–E4923. 10.1073/pnas.1704553114.

37. Zhao, B., Socha, J., Toth, A., Fernandes, S., Warheit-Niemi, H., Ruff, B., Khurana Hershey, G.K., VanDussen, K.L., Swarr, D., and Zacharias, W.J. (2024). The Homeobox Transcription Factor Cux1 Coordinates Postnatal Epithelial Developmental Timing but Is Dispensable for Lung Organogenesis and Regeneration. American Journal of Respiratory Cell and Molecular Biology. 10.1165/rcmb.2024-0147OC.

38. Chang, L., Azzolin, L., Di Biagio, D., Zanconato, F., Battilana, G., Lucon Xiccato, R., Aragona, M., Giulitti, S., Panciera, T., Gandin, A., et al. (2018). The SWI/SNF complex is a mechanoregulated inhibitor of YAP and TAZ. Nature 563, 265–269. 10.1038/s41586-018-0658-1.

39. Gokey, J.J., Snowball, J., Sridharan, A., Sudha, P., Kitzmiller, J.A., Xu, Y., and Whitsett, J.A. (2021). YAP regulates alveolar epithelial cell differentiation and AGER via NFIB/KLF5/NKX2-1. iScience 24, 102967. 10.1016/j.isci.2021.102967.

40. Frank, D.B., Peng, T., Zepp, J.A., Snitow, M., Vincent, T.L., Penkala, I.J., Cui, Z., Herriges, M.J., Morley, M.P., Zhou, S., et al. (2016). Emergence of a Wave of Wnt Signaling that Regulates Lung Alveologenesis by Controlling Epithelial Self-Renewal and Differentiation. Cell Reports 17, 2312–2325. 10.1016/j.celrep.2016.11.001.

41. Riou, R., Ladli, M., Gerbal-Chaloin, S., Bossard, P., Gougelet, A., Godard, C., Loesch, R., Lagoutte, I., Lager, F., Calderaro, J., et al. (2020). ARID1A loss in adult hepatocytes activates β-catenin-mediated erythropoietin transcription. eLife 9, e53550. 10.7554/eLife.53550.

42. Ellis, L.V., Cain, M.P., Hutchison, V., Flodby, P., Crandall, E.D., Borok, Z., Zhou, B., Ostrin, E.J., Wythe, J.D., and Chen, J. (2020). Epithelial Vegfa Specifies a Distinct Endothelial Population in the Mouse Lung. Developmental Cell 52, 617-630.e616. 10.1016/j.devcel.2020.01.009.

43. Toth, A., Kannan, P., Snowball, J., Kofron, M., Wayman, J.A., Bridges, J.P., Miraldi, E.R., Swarr, D., and Zacharias, W.J. (2023). Alveolar epithelial progenitor cells require Nkx2-1 to maintain progenitor-specific epigenomic state during lung homeostasis and regeneration. Nature Communications 14, 8452. 10.1038/s41467-023-44184-0.

44. Peng, D., Si, D., Zhang, R., Liu, J., Gou, H., Xia, Y., Tian, D., Dai, J., Yang, K., Liu, E., et al. (2017). Deletion of SMARCA4 impairs alveolar epithelial type II cells proliferation and aggravates pulmonary fibrosis in mice. Genes & Diseases 4, 204–214. 10.1016/j.gendis.2017.10.001.

45. Glaros, S., Cirrincione, G.M., Palanca, A., Metzger, D., and Reisman, D. (2008). Targeted Knockout of BRG1 Potentiates Lung Cancer Development. Cancer Research 68, 3689–3696. 10.1158/0008-5472.CAN-07-6652.

46. Orvis, T., Hepperla, A., Walter, V., Song, S., Simon, J., Parker, J., Wilkerson, M.D., Desai, N., Major, M.B., Hayes, D.N., et al. (2014). BRG1/SMARCA4 Inactivation Promotes Non–Small Cell Lung Cancer Aggressiveness by Altering Chromatin Organization. Cancer Research 74, 6486–6498. 10.1158/0008-5472.CAN-14-0061.

47. Concepcion, C.P., Ma, S., LaFave, L.M., Bhutkar, A., Liu, M., DeAngelo, L.P., Kim, J.Y., Del Priore, I., Schoenfeld, A.J., Miller, M., et al. (2022). Smarca4 Inactivation Promotes Lineage-Specific Transformation and Early Metastatic Features in the Lung. Cancer Discovery 12, 562–585. 10.1158/2159-8290.CD-21-0248.

48. Schuettengruber, B., Bourbon, H.-M., Croce, L.D., and Cavalli, G. (2017). Genome Regulation by Polycomb and Trithorax: 70 Years and Counting. Cell 171, 34–57. 10.1016/j.cell.2017.08.002.

49. Snitow, M.E., Li, S., Morley, M.P., Rathi, K., Lu, M.M., Kadzik, R.S., Stewart, K.M., and Morrisey, E.E. (2015). Ezh2 represses the basal cell lineage during lung endoderm development. Development 142, 108–117. 10.1242/dev.116947.

50. Ferrari, K.J., Scelfo, A., Jammula, S., Cuomo, A., Barozzi, I., Stützer, A., Fischle, W., Bonaldi, T., and Pasini, D. (2014). Polycomb-Dependent H3K27me1 and H3K27me2 Regulate Active Transcription and Enhancer Fidelity. Molecular Cell 53, 49–62. 10.1016/j.molcel.2013.10.030.

51. Liberti, D.C., Wen, H., Quansah, K.K., Chandrasekaran, P., Pankin, J., Michki, N.S., Jin, A., Lu, M., Nieuwburgh, M.P.D., Young, L.R., et al. (2023). Endodermal BRD4 mediates epithelial-mesenchymal crosstalk during lung development. 10.1101/2023.05.21.541621.

52. Devaiah, B.N., Case-Borden, C., Gegonne, A., Hsu, C.H., Chen, Q., Meerzaman, D., Dey, A., Ozato, K., and Singer, D.S. (2016). BRD4 is a histone acetyltransferase that evicts nucleosomes from chromatin. Nat Struct Mol Biol 23, 540–548. 10.1038/nsmb.3228.

53. Du, J., Liu, Y., Sun, J., Yao, E., Xu, J., Wu, X., Xu, L., Zhou, M., Yang, G., and Jiang, X. (2024). ARID1A safeguards the canalization of the cell fate decision during osteoclastogenesis. Nature Communications 15, 5994. 10.1038/s41467-024-50225-z.

54. Dannappel, M.V., Zhu, D., Sun, X., Chua, H.K., Poppelaars, M., Suehiro, M., Khadka, S., Sian, T.C.C.L.K., Sooraj, D., Loi, M., et al. (2022). CDK8 and CDK19 regulate intestinal differentiation and homeostasis via the chromatin remodeling complex SWI/SNF. The Journal of Clinical Investigation 132. 10.1172/JCI158593.

55. He, H., Bell, S.M., Davis, A.K., Zhao, S., Sridharan, A., Na, C.-L., Guo, M., Xu, Y., Snowball, J., Swarr, D.T., et al. (2024). PRDM3/16 regulate chromatin accessibility required for NKX2-1 mediated alveolar epithelial differentiation and function. Nature - Communications 15, 8112. 10.1038/s41467-024-52154

56. Frank, D.B., Penkala, I.J., Zepp, J.A., Sivakumar, A., Linares-Saldana, R., Zacharias, W.J., Stolz, K.G., Pankin, J., Lu, M., Wang, Q., et al. (2019). Early lineage specification defines alveolar epithelial ontogeny in the murine lung. Proceedings of the National Academy of Sciences 116, 4362–4371. 10.1073/pnas.1813952116.

57. Zacharias, W.J., Frank, D.B., Zepp, J.A., Morley, M.P., Alkhaleel, F.A., Kong, J., Zhou, S., Cantu, E., and Morrisey, E.E. (2018). Regeneration of the lung alveolus by an evolutionarily conserved epithelial progenitor. Nature 555, 251–255. 10.1038/nature25786.

58. Sinner, D.I., Carey, B., Zgherea, D., Kaufman, K.M., Leesman, L., Wood, R.E., Rutter, M.J., de Alarcon, A., Elluru, R.G., Harley, J.B., et al. (2019). Complete Tracheal Ring Deformity. A Translational Genomics Approach to Pathogenesis. American Journal of Respiratory and Critical Care Medicine 200, 1267–1281. 10.1164/rccm.201809-1626OC.

59. McGinnis, C.S., Murrow, L.M., and Gartner, Z.J. (2019). DoubletFinder: Doublet Detection in Single-Cell RNA Sequencing Data Using Artificial Nearest Neighbors. Cell Syst 8, 329-337.e324. 10.1016/j.cels.2019.03.003 PMID - 30954475.

60. Mohammed, H., Taylor, C., Brown, G.D., Papachristou, E.K., Carroll, J.S., and D’Santos, C.S. (2016). Rapid immunoprecipitation mass spectrometry of endogenous proteins (RIME) for analysis of chromatin complexes. Nature Protocols 11, 316–326. 10.1038/nprot.2016.020.

